# Widespread distribution of collagens and collagen-associated domains in eukaryotes

**DOI:** 10.1101/2021.10.08.463732

**Authors:** Tess A. Linden, Nicole King

## Abstract

The origin of collagen, the dominant structural component of metazoan extracellular matrix, has long been cited as a critical step in the evolution of metazoan multicellularity. While collagens were once thought to be found only in metazoans, scattered reports of collagen domains in Fungi, and more recently in close relatives of metazoans, have called into question whether collagens are truly unique to metazoans. Here, we take advantage of recently sequenced genomes and transcriptomes of diverse holozoans (the clade encompassing metazoans and their close relatives), as well as publicly available proteomes from diverse non-holozoan eukaryotes, to conduct a systematic search for collagen domains across eukaryotic diversity. We find that collagen domains are ubiquitous in choanoflagellates, the sister group of metazoans, and widespread across many other major eukaryotic taxa. Many predicted collagens in non-metazoans are comparable to metazoan collagens in length and proline content. Moreover, most are present in species that also encode putative prolyl 4-hydroxylase domains, suggesting that, like metazoan collagens, they may be stabilized through the hydroxylation of prolines. Fibrillar collagen and collagen IV appear to be unique to metazoans, and we posit that their ability to assemble into superstructures may have contributed to the origin of metazoan multicellularity.

## Introduction

In metazoans, the extracellular matrix (ECM) is indispensable to development, form, and function. Collagen is the major structural protein of metazoan ECM (Frantz et al., 2010), and is the most abundant protein in metazoan bodies by mass, making up one third of the dry weight of an adult mammal (Shoulders and Raines, 2009) and over 70% of the weight of some sponges (Ehrlich et al., 2018). The diagnostic feature of collagens is the repeated triplet motif Gly-X-Y, in which X and Y can be any amino acid (Box 1). The repetition of the collagen repeat motif Gly-X-Y leads to the formation of a stable, rod-like triple helix, which in turn supplies tensile strength to metazoan tissues (Buehler, 2006; Shoulders and Raines, 2009; Wenger et al., 2007).

For most of the twentieth century, collagen was thought to be unique to metazoans and absent from other lineages with complex multicellularity (e.g., plants and fungi). Thus, because of its apparent phylogenetic restriction and its importance to metazoan ECM, collagen was repeatedly hypothesized to be one of the key innovations that allowed metazoans and their characteristic tissues to evolve (Erwin, 1993; Towe, 1970). The hypothesized connection between metazoan origins and the evolution of collagen was brought into question with the sequencing of genomes from three close relatives of metazoans—two choanoflagellates and one filasterean—each of which encode collagen domains (Fairclough et al., 2013; Grau-Bové et al., 2017; Hynes, 2012; King et al., 2008). These joined reports of putative collagens in Dikarya Fungi (Celerin et al., 1996; de Bruin et al., 2002; Wang and Leger, 2006), the malaria parasite *Plasmodium falciparum* (Rasmussen et al., 2003), and the amoebozoan *Dictyostelium discoideum* (Fidler et al., 2017) to create a growing catalog of putative collagens in non-metazoan eukaryotes. In addition, collagen domains have been detected in several species of bacteria (Bachert et al., 2015; Charalambous et al., 1988; Ghosh et al., 2012; Kananavičiūtė et al., 2020; Lukomski et al., 2017; Mohs et al., 2007; Rasmussen et al., 2003; Yu et al., 2014) and viruses (Bamford and Bamford, 1990; La Scola et al., 2008; Luther et al., 2011; Medveczky et al., 1993; Raoult et al., 2004; Tidona and Darai, 1997; van Hulten et al., 2001; Yau et al., 2011; Zhang et al., 2004), although due to their evolutionary distance, these were typically hypothesized to have evolved independently from metazoan collagens (Rasmussen et al., 2003) or through horizontal gene transfer from metazoans (Luther et al., 2011; Rasmussen et al., 2003). Nonetheless, the growing number of reports has pointed to the possibility that collagens might be more ancient and/or more widespread than previously realized.

Therefore, to better understand the phylogenetic distribution and ancestry of collagen domains across the tree of eukaryotes, we analyzed recently sequenced genomes and transcriptomes of close relatives of metazoans, the choanoflagellates and other non-metazoan holozoans (Table S1), as well as those of diverse other eukaryotes representing many major clades. We found that collagen domains are common in the close relatives of metazoans, including choanoflagellates, many of which encode extended collagen domains comparable in length and proline content to metazoan collagens. Furthermore, we found that collagen domains are widely distributed across many eukaryotes that are only distantly related to metazoans, including Fungi, Archaeplastida, and members of the SAR clade. In contrast, canonical metazoan collagens such as fibrillar collagens and collagen IV are apparently restricted to metazoans and may help explain unique features of metazoan biology.

### Box 1.

**Collagen terminology.**

The criteria for what constitutes “a collagen” are not well defined (Garrone, 1999; Gay and Miller, 1983; Ricard-Blum, 2011). The major defining characteristic of a collagen is the collagen triple helix, but some authors restrict the term “a collagen” to mean a protein that plays a structural role in the ECM (Garrone, 1999; Gay and Miller, 1983), while others define “a collagen” as any protein containing a collagen triple helix, including intracellular proteins of the immune system (Casals et al., 2019; Fraser and Tenner, 2008) or transmembrane signaling proteins (Maertens et al., 2007). Here, we adopt the latter usage, for two reasons: first, many of the proteins we discuss have not been functionally characterized, and whether a given protein plays a structural role in the ECM is difficult to predict from sequence alone. Second, this usage acknowledges the possibility that collagens could evolve dynamically between structural and non-structural roles or between intracellular and extracellular localizations over evolutionary time. For brevity and simplicity, we use the term “collagen domain” to refer to any stretch of collagen triple helix repeats, i.e., (Gly-X-Y)_n_.

## Results

### Collagen domains and conserved collagen-associated domains predate metazoan origins

To start, we focused our search on close relatives of metazoans, hypothesizing that they were most likely to illuminate the proximal ancestry of metazoan collagens (Fig. 1). From the 29 genomes and transcriptomes analyzed (Table S1), we found that all 22 choanoflagellates, three of four filastereans (*Ministeria vibrans, Pigoraptor vietnamica*, and *Pigoraptor chileana)* and one of two ichthyosporeans (*Sphaeroforma arctica*) encoded proteins with collagen domains (domains annotated by InterProScan as IPR008160). Like metazoan collagens, many of these proteins were predicted to carry transmembrane regions and/or N-terminal signal sequences (Fig. S1), suggesting that they are secreted into the ECM. Furthermore, all 29 species were found to encode putative prolyl 4-hydroxylase domains (IPR006620), the class of enzyme responsible for the post-translational modification of proline to 4-hydroxyproline, a step thought to be required for collagen triple helix formation in metazoans (Berg and Prockop, 1973; Canty and Kadler, 2005; Juva et al., 1966; Kao et al., 1979; Walmsley et al., 1999). (The strictness of this requirement has been challenged by observations of triple helix formation without hydroxyproline in some contexts, such as hymenopteran cocoons (Sutherland et al., 2013), recombinant non-metazoan cells (Olsen et al., 2001; Perret et al., 2001; Ruggiero et al., 2000), and bacteria (Yu et al., 2014).) Thus, 26 of the 29 non-metazoan holozoans that we analyzed appear to have the genetic machinery required to produce collagen triple helices that may be modified with 4-hydroxyproline.

**Figure 1.**
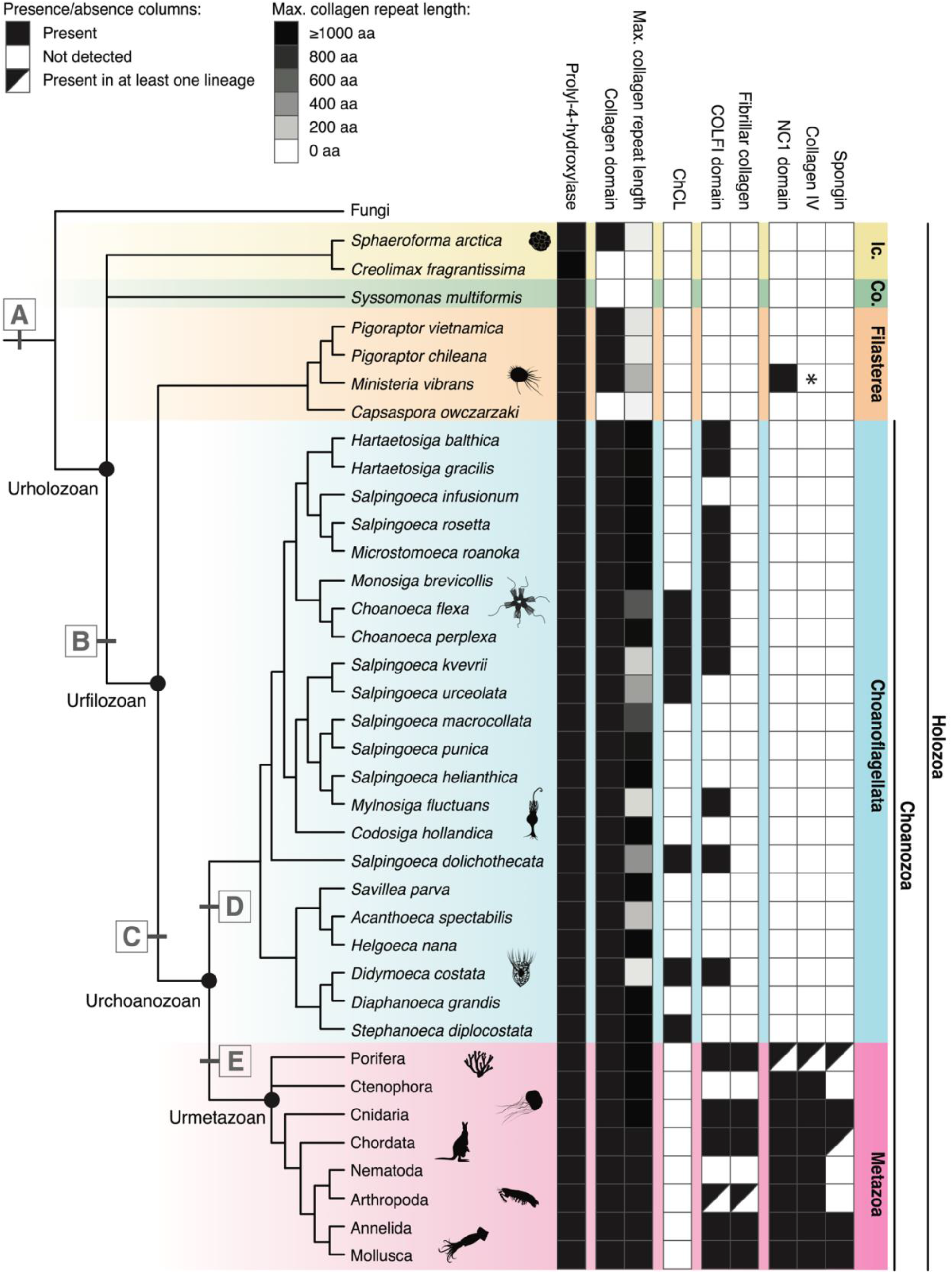
Collagen domains and collagen-associated domains are conserved in close relatives of Metazoa. We detected collagen domains (IPR008160) and putative prolyl 4-hydroxylase domains (IPR006620) in the predicted proteomes of all metazoans and choanoflagellates and in most other holozoans analyzed, and found that the distributions of other collagen-associated domains such as COLF1 (IPR000885) and NC1 (IPR001442/IPR036954) are also consistent with pre-metazoan origins. Holozoans include choanoflagellates, filastereans, corallochytreans (“Co.”) and ichthyosporeans (“Ic.”). Collagen length in choanoflagellates rivals that of metazoans, as shown here in the “Max. collagen repeat length” column, in which the maximum collagen repeat length of any protein detected in the species’ predicted proteome is indicated by degree of shading (see key and Methods). **(A – E)** Mapping the distribution of collagen domains and key collagen-associated domains helps resolve the pre-metazoan evolution of collagens. **(A)** Putative prolyl 4-hydroxylase domains, which in metazoans perform a post-translational modification step important for collagen triple helix formation, were detected in all 29 non-metazoan holozoans. This domain is also present in many non-holozoan eukaryotes (Fig. S6), and thus might predate the divergence between Fungi and Holozoa. **(B)** The NC1 domain characteristic of metazoan collagen IV was detected in a collagen domain-containing protein (MvCN) of the filasterean *Ministeria vibrans* as previously reported (Grau-Bové et al., 2017), but not in any other holozoans, including choanoflagellates. Thus, the NC1 domain was likely present in the common ancestor of filastereans and choanozoans. **(C)** COLF1 domains, which are diagnostic of metazoan fibrillar collagens, are broadly distributed across choanoflagellates and metazoans, indicating that they predate the Urchoanozoan. However, in choanoflagellates, the COLF1 domain was not detected in any proteins that also contained collagen domains. **(D)** ChCL, a choanoflagellate-specific transmembrane protein consisting of a collagen domain and a lectin/glucanase domain, is broadly distributed across the choanoflagellate phylogeny, suggesting that it was likely present in the last common ancestor of choanoflagellates. **(E)** Fibrillar collagens (those containing COLF1 and collagen domains) appear to be restricted to metazoans. Likewise, canonical collagen IVs (proteins with >100 Gly-X-Y repeats and a C-terminal NC1 domain with a conserved HSQ motif) and spongins (truncated collagen IV variants lacking the conserved HSQ motif) are also restricted to metazoans. *: The MvCN protein was previously described as a collagen IV (Grau-Bové et al., 2017), but given the differences between MvCN and metazoan collagen IV, it is currently unclear whether this filasterean protein is homologous to collagen IV (see main text). Figure format is adapted with permission from Figure 1 of Brunet et al. (2017). Consensus phylogeny of choanoflagellate relationships is based on Richter et al. (2018).

At least two classes of collagens may have been present in the Urmetazoan (the last common ancestor of all metazoans) based on their broad phylogenetic distribution within metazoans. The first is fibrillar collagen, which is characterized by a long, uninterrupted collagen domain and a C-terminal COLF1 domain (Exposito et al., 2010). The second is collagen IV, which is characterized by a triple helix with many interruptions and a C-terminal NC1 domain (Boute et al., 1997; Exposito et al., 2002; Fidler et al., 2018, 2017). Included in this second class is the collagen IV variant spongin, which is distinguished by a truncated collagen domain and the lack of a conserved HSQ motif in its NC1 domain (Aouacheria et al., 2006; Fidler et al., 2017). To determine whether these key metazoan collagens may have predated the Urmetazoan, we searched our data set for proteins with collagen domains and COLF1 or NC1 domains. Prior investigations found that *S. rosetta* and *M. brevicollis* encode COLF1 domains homologous to the C-terminal domains of metazoan fibrillar collagens, but that these domains are not present in the same proteins as collagen domains (Hynes, 2012; King et al., 2008). We found that this pattern holds true across choanoflagellate diversity: COLF1 was detected in 12 of 22 choanoflagellate proteomes (Fig. 1), but none of the choanoflagellate COLF1 domains were present in proteins with collagen domains. COLF1 was not detected in any other holozoans. Thus, the COLF1 domain likely evolved on the Choanozoan stem lineage, but canonical fibrillar collagens appear to have evolved, presumably through domain shuffling, after the split between the choanoflagellate and metazoan lineages.

Next, we analyzed whether any non-metazoan holozoans encoded NC1 domains. Consistent with previous findings in *S. rosetta* and *M. brevicollis* (Fidler et al., 2017), we did not detect NC1 domains in any of the 22 choanoflagellate proteomes. Outside choanozoans, we detected an NC1 domain only in one protein encoded by the filasterean *Ministeria vibrans* (Fig. 1; Fig. S2C), as previously reported by Grau-Bové et al. (2017). Like metazoan collagen IV, this *M. vibrans* protein (hereafter “MvCN”, for *Ministeria vibrans* <u>C</u>ollagen + <u>N</u>C1) contains collagen domains and a C-terminal NC1 domain (Fig. S2C). This raises the question of whether MvCN might be homologous to collagen IV, and thus that collagen IV might be an ancient protein that predates the last common ancestor of filastereans and metazoans. To investigate this possibility more closely, we examined each of the domains of MvCN individually. By aligning the MvCN NC1 domain with metazoan collagen IVs, we found that it contains the conserved HSQ motif found in the NC1 domains of metazoan collagen IVs but not in those of spongins (Fig. S3B) (Fidler et al., 2017). Next, we examined the MvCN collagenous region. In contrast with metazoan collagen IV, the MvCN collagenous region is much shorter: it contains a total of 81 Gly-X-Y repeats, as opposed to the hundreds of Gly-X-Y repeats typical of metazoan collagen IV: e.g., 443 repeats in *Mus musculus* (UniProt ID P02463), 264 repeats in the mollusk *Lottia gigantea* (UniProt ID V4A1B9), or 473 repeats in the cnidarian *Nematostella vectensis* (UniProt ID V9GW22). Finally, we examined its non-collagenous domain. Unlike metazoan collagen IV, MvCN contains a long, N-terminal non-collagenous domain of unknown function. Through BLAST search, we found that this domain shares sequence similarity with uncharacterized bacterial proteins (Fig. S3A), indicating that MvCN may be a eukaryotic-bacterial fusion protein. Thus, the NC1 domain appears to predate the divergence between filastereans and choanozoans, but whether MvCN shares a common ancestry with metazoan collagen IV/spongin remains unclear. It is possible, for example, that an NC1 domain was present in a protein without collagen domains in a stem holozoan and became fused to collagen domains independently in the filasterean and metazoan lineages.

### Choanoflagellates encode collagens with diverse non-collagenous domains

Because choanoflagellates are the closest living relatives of metazoans and provide a unique window into the ancestry of metazoan genes, we examined the protein domain architecture of their collagens in more detail. The non-collagenous protein domain found most frequently in choanoflagellate collagens was the von Willebrand Factor A (VWFA) domain, a domain commonly found in metazoan ECM proteins such as collagen VI and FACIT collagens (Fig. S2D) (Ricard-Blum, 2011). VWFA domains often mediate protein-protein interactions, facilitating the binding of ECM proteins to each other (Hynes and Naba, 2012).

Interestingly, two choanoflagellate species encode collagens with domains that are also present in metazoan collagenous immune proteins. The choanoflagellate *Salpingoeca kvevrii* encodes a collagen with an SRCR domain, a combination found in some metazoan scavenger receptors that play a role in phagocytosis and innate immunity (Fig. S2D) (Gowen et al., 2001; Kodama et al., 1996; Neubauer et al., 2016; Yamada et al., 1998). *Microstomoeca roanoka* encodes a collagenous protein with a gamma fibrinogen domain, making it similar to ficolins, a group of collagens that function in innate immune system activation in chordates (Endo et al., 2015; Matsushita, 2010). Because both species nest deeply within choanoflagellates, we posit that these choanoflagellate proteins evolved independently from metazoan scavenger receptors and ficolins. Nonetheless, the shared combination of protein domains raises the possibility that these collagens might play analogous roles in bacterial recognition, phagocytosis, or immunity in choanoflagellates in a manner similar to the metazoan defense collagens.

Because the choanoflagellate clade contains as much genetic diversity as its sister group, the Metazoa (Richter et al., 2018), we were interested in whether our data set could reveal patterns of collagen evolution within the choanoflagellate lineage. The diversity of collagen repertoires across choanoflagellates and the high degree of variability in collagen number and domain structure between closely related species suggested that collagens are rapidly evolving within choanoflagellates. For example, *Choanoeca flexa* was found to encode six collagens, whereas its closely related sister species *C. perplexa* encodes 16 collagens, and their next-closest relative, *M. brevicollis*, encodes only two (Fig. S1); many of these proteins have unique domain structures without clear orthologs in the other species. By contrast, some collagens appear to be conserved across choanoflagellate evolution. Mirroring the conserved collagens of metazoans (collagen IV and fibrillar collagens), we identified a possibly ancient collagen that is distributed widely within choanoflagellates. This protein, which we refer to as “ChCL” (for <u>Ch</u>oanoflagellate <u>C</u>ollagenous <u>L</u>ectin) has a characteristic architecture of a ∼20-amino acid N-terminal transmembrane domain followed by an extracellular region with a ∼50-amino acid collagen domain, a ∼700-amino acid non-collagenous domain, and a ∼200-amino acid C-terminal Concanavalin A-like lectin/glucanase domain (IPR013320) (Fig. S4A). The non-collagenous domain of ChCL has sequence similarity with non-collagenous metazoan proteins of unknown function (Fig. S4B), while the C-terminal domain closely resembles genes of bacterial origin (such as MSX41286.1, an Actinobacterium protein of unknown function, or WP_007139689.1, a putative cell wall-associated protein in a Flavobacterium) (Fig. S4C). Based on its distribution and on the most recent understanding of choanoflagellate phylogenetic relationships, ChCL appears to have been present in the last common ancestor of all choanoflagellates (Fig. 1).

### Collagen domains are widespread across eukaryotes

The finding that collagenous proteins are nearly ubiquitous in the closest living relatives of metazoans inspired us to broaden our search to include more distant lineages. To investigate the distribution of collagen domains across eukaryotic diversity, we took advantage of the proteomes available in the UniProtKB database (UniProt Consortium, 2021), which includes representatives of 288 families of non-metazoan eukaryotes across many, but not all, of the major eukaryotic groups (Fig. 2A). We detected proteins with collagen domains in nearly all major eukaryotic groups analyzed (Fig. 2B). For example, out of 169 fungal families in our analysis, 73 families encode at least one protein with a collagen domain (IPR008160) (Fig. 2C). Families encoding collagens were concentrated within Ascomycota, but we also detected collagens sparsely distributed in Basidiomycota, Zoopagomycota, Mucoromycota, Cryptomycota, and Microsporidia (Fig. S5B). Within Archaeplastida, 17 out of 71 families encode collagens (Fig. 2D). Nine of these 17 families are within land plants (Embryophyta), revealing that collagens are present in at least two lineages with complex multicellularity. In the SAR clade, 19 out of 31 families were found to encode collagens (Fig. 2E). The only major eukaryotic group in which we detected no collagen domains was Haptista (Fig. 2F), which is represented by only two proteomes in our data set.

**Figure 2.**
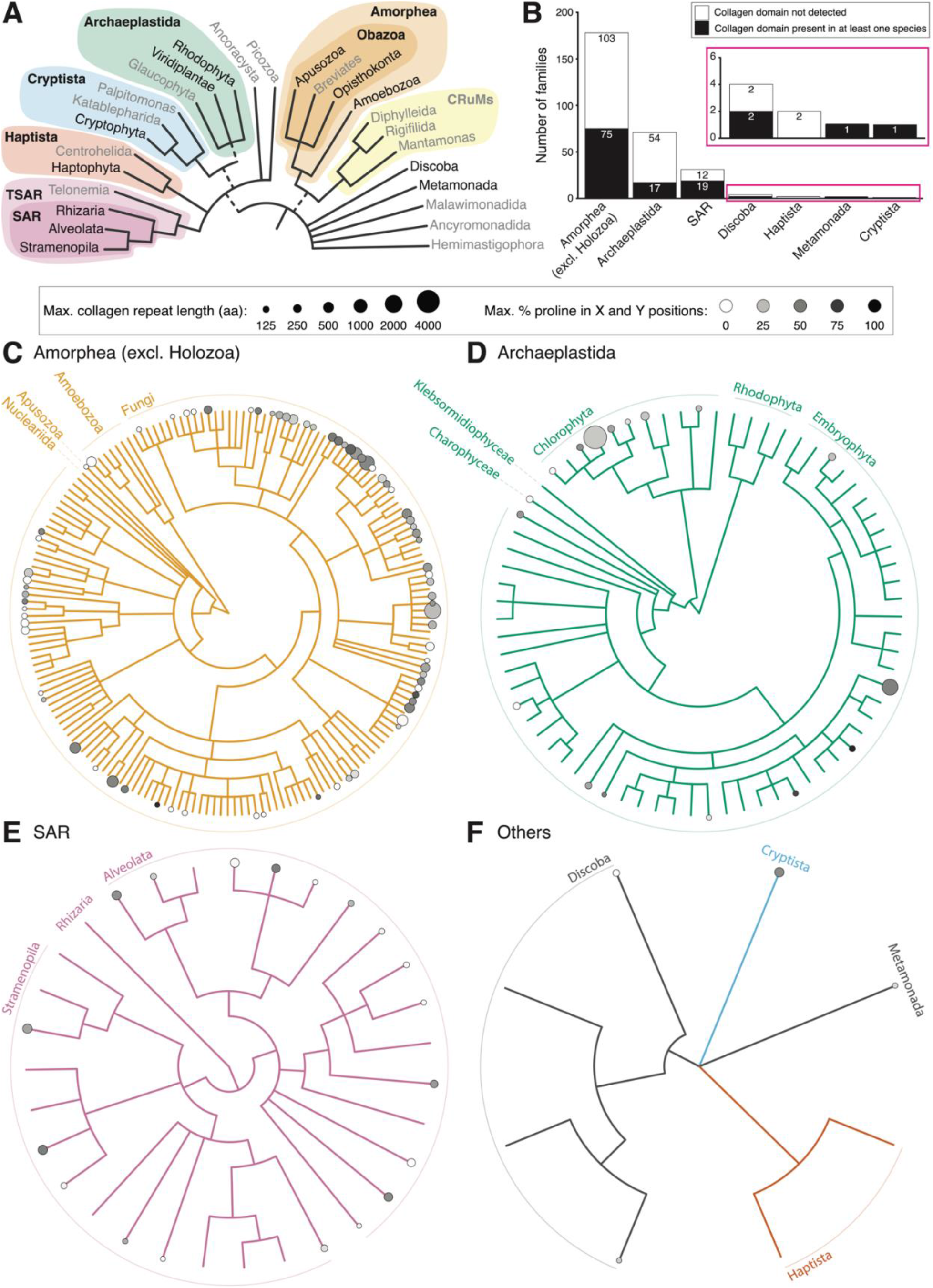
Collagen domains are detected throughout eukaryotic diversity. Having found that collagens are widespread in holozoans, we broadened our search to investigate whether collagen domains are found in other eukaryotic taxa. Our analysis included 2,598 eukaryotic proteomes present in the UniProtKB database that met quality criteria based on BUSCO score and protein count (see Methods). **(A)** The current consensus eukaryotic tree. The UniProtKB database contains proteomes from many branches of the eukaryotic tree of life, but not all. Clades shown in black are represented by at least one proteome in UniProtKB, while clades in gray are unrepresented. Dashed lines represent uncertainty about the monophyly of the indicated groups. **(B)** For each major eukaryotic group, the number of analyzed families in which a collagen domain was detected (black bar) or not detected (white bar) is shown (y-axis). For the Amorphea supergroup, which contains the Holozoa, we only show data for non-holozoans. Collagen domains were detected in at least one family from all eukaryotic groups analyzed, with the exception of Haptista. Even when omitting Holozoa, proteomes from Amorphea (particularly Fungi; see panel C) are heavily overrepresented in the UniProtKB database. Meanwhile, several major eukaryotic groups are represented by <10 families each (inset). **(C** to **F)** Collagen domains are broadly distributed not just across, but within, all major eukaryotic groups. In these trees, each stem represents one taxonomic family for which which at least one proteome meeting our quality criteria is present in the UniProtKB database. The presence of a circle indicates that at least one species in that family encodes a collagen domain (IPR008160). The size of the circle represents the maximum collagen repeat length of any collagen in the family, while the shading of the circle represents the maximum proline content of any collagen in the family (see Methods). Note that the maximum collagen repeat length and maximum proline content may be from different proteins. **(C)** Distribution of collagen domains across Amorphea families. Holozoans were excluded in order to highlight that collagen domains are broadly distributed even in distant relatives of metazoans. **(D)** Distribution of collagen domains across Archaeplastida families. The chlorophyte *Coccomyxa subellipsoidea* encodes the longest collagen domain detected in our entire data set, with a collagen repeat length of >4000 amino acids. **(E)** Distribution of collagen domains across SAR families. Collagen domains were detected in Stramenopiles and Alveolates, but not Rhizarians. **(F)** Distribution of collagen domains across the families of the other major groups of eukaryotes: Haptista, Cryptista, Metamonada, and Discoba. Haptista, which is represented by only two families in our analysis, is the only eukaryotic group for which no collagen domains were detected. Panel A was adapted with permission from Figure 1 of Burki et al. (2020).

The low information-density of the collagen domain (Box 1) makes it difficult to infer whether the collagen domains found in diverse eukaryotic clades are homologous with metazoan collagens or whether they evolved convergently. However, we can ask if non-metazoan collagen domains resemble metazoan collagens in length and proline content, and whether the collagen-encoding species also encode a putative prolyl 4-hydroxylase. We found that most non-metazoan eukaryotic proteomes in our data set encode at least one protein with a putative prolyl 4-hydroxylase domain (Fig. S6); specifically, 84% of the 2,011 proteomes that encode collagens also encode a putative prolyl 4-hydroxylase. Whether an enzyme is capable of hydroxylating proline in a collagen-like substrate is difficult to predict from sequence alone, but such enzymatic activity outside metazoans would not be unprecedented: endogenous enzymes with collagen prolyl 4-hydroxylase activity have been reported in plants (Hieta and Myllyharju, 2002) and Fungi (de Bruin et al., 2002). Next, we compared the collagen repeat length across all collagens detected in metazoans (*n* = 57,104 collagens), choanoflagellates (*n* = 258 collagens), Archaeplastida (*n* = 75 collagens), Fungi (*n* = 508 collagens), and SAR (*n* = 176 collagens) (Fig. 3A; Fig. S7A). Metazoan collagens are the longest, with a median collagen repeat length of 192 amino acids (i.e., a median of 64 Gly-X-Y repeats per protein); choanoflagellate collagens have the second-longest, with a median collagen repeat length of 145.5 amino acids; and SAR collagens have the shortest, with a median of 85.5 amino acids. We then quantified proline content across the same data set (Fig. 3B; Fig. S7B) and found that metazoan collagen domains contain the most proline at the X and Y amino acid positions, with a median of 29.9% proline. Choanoflagellate collagens were the second-most proline-rich at 15.6%, while SAR collagens were the least proline-rich at a median of 0%. While the median metazoan collagen may have longer collagen domains and more proline than the median non-metazoan collagen, it is notable that the distributions of collagen repeat length and proline content overlap significantly across all these eukaryotic groups (Fig. 3). Together, these data suggest that some non-metazoans likely express collagen triple helices with similar structural qualities to metazoan collagens, which could therefore play similar roles in their biology.

**Figure 3.**
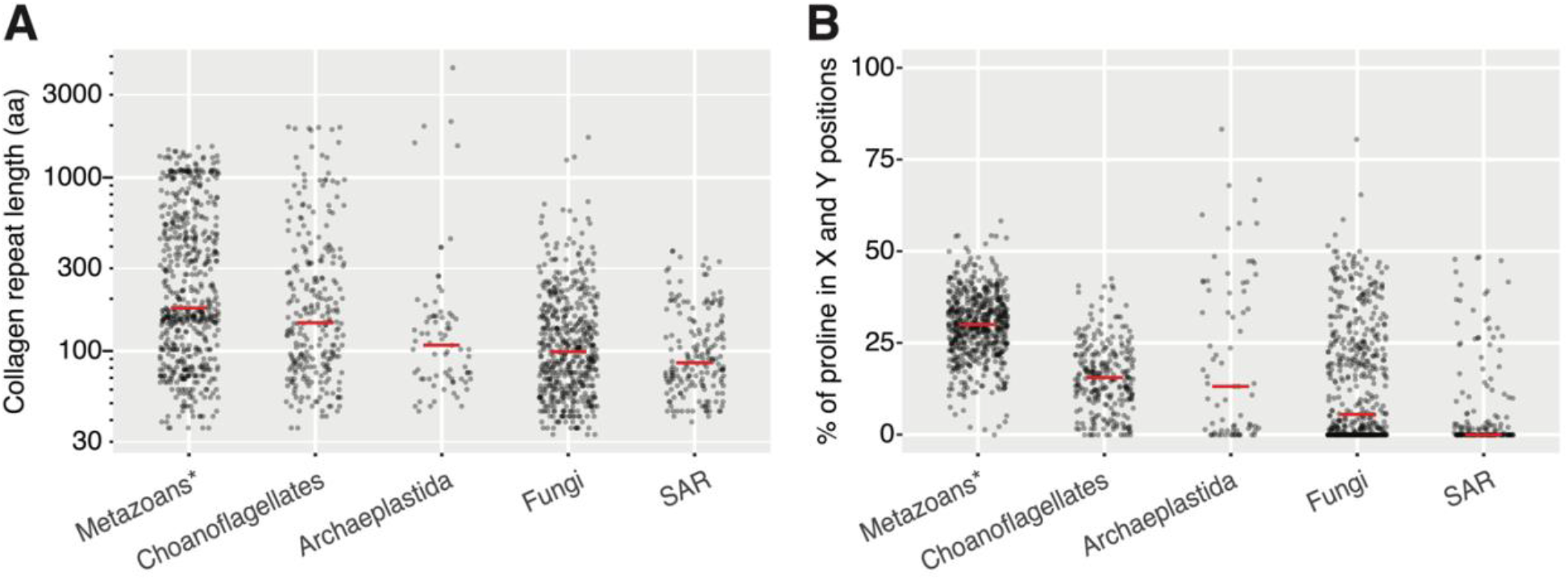
Metazoans encode collagens with more collagen repeats and higher proline content than other eukaryotes. To compare the characteristics of collagen domains across eukaryotic taxa, we examined collagen repeat length and proline content (see Methods) across all collagens detected in all analyzed species of choanoflagellates (*n* = 258 collagens), metazoans (*n =* 57,104 collagens), Archaeplastida (*n* = 75 collagens); fungi (*n* = 508 collagens); and SAR (*n* = 176 collagens). **(A)** Metazoans have the highest total collagen repeat length per protein (median = 192 amino acids), choanoflagellates the second-highest (median = 145.5 amino acids), and SAR the lowest (median = 85.5 amino acids). Note that the y-axis is on a log-scale. **(B)** Metazoan collagens have the highest proline content (median = 29.9%), choanoflagellates have the second-highest (median = 15.6%), and SAR the lowest (median = 0%). Thus, metazoan collagens are characterized by a greater number of repeats and higher proline content compared to those of other eukaryotic taxa; however, it is notable that the distributions of these characteristics across all five taxa are highly overlapping. Red bars represent medians. *: to more clearly visualize the distribution of the metazoan collagen data, a randomly selected 1% of metazoan collagens are shown here. For comparison of collagen repeat length and proline content across all eukaryotic groups analyzed, see Figure S7.

## Discussion

Extracellular proline-rich glycoproteins have long been known to exist in non-metazoan eukaryotes (Davies et al., 1997; Hallmann, 2006; Lamport, 1977; Perfect et al., 1998; Showalter, 1993; Sumper and Hallmann, 1998). However, collagen domains have rarely been reported in distant eukaryotic relatives of Metazoa. The furthest examples, to our knowledge, have been reports of collagen repeats in the genomes of the apicomplexan (SAR) parasite *Plasmodium falciparum* (Rasmussen et al., 2003) and the amoebozoan *Dictyostelium discoideum* (Fidler et al., 2017), and of cell-surface proteins with collagen repeats in a handful of species of Dikarya fungi (Celerin et al., 1996; de Bruin et al., 2002; Wang and Leger, 2006). Our findings suggest that collagen domains may be a near-ubiquitous feature of holozoans (Fig. 1, Fig. S5C) and may also be widespread across eukaryotic diversity (Fig. 2). The collagen domain is characterized by a repeated triplet motif (Box 1) whose simplicity makes it a reasonable candidate for repeated convergent evolution (Chen et al., 1997; Stern, 2013). With the exception of metazoans (Fig. S5A), choanoflagellates (Fig. S5C), and perhaps ascomycete Fungi (Fig. S5B), the distribution of collagen domains across all of the eukaryotic groups in our data set is notably scattered: many pairs of sister families differ from each other in whether they encode collagen domains (Fig. 2C-F). Such a pattern could indicate either rampant convergent evolution or rampant loss. Due to the inherent difficulty of aligning highly repetitive domains, it remains unclear how many times collagen domains may have evolved in the history of eukaryotes. Even if the collagen domains outside Metazoa evolved independently from those within Metazoa or Holozoa, it is nonetheless possible that proteins with collagen-like properties might be important to the biology of many organisms across eukaryotic diversity, not just metazoans.

The evolution of collagen has often been cited as a key prerequisite for metazoan complex multicellularity (Erwin, 1993; Fidler et al., 2018; Reinhard et al., 2016; Towe, 1970). Here, we found that while collagen domains are encoded by diverse eukaryotes, several features distinguish metazoan collagen repertoires from those of non-metazoans. Metazoan collagen domains tend to be slightly longer and more proline-rich than those of any other eukaryotic clade in our data set (Fig. 3). Furthermore, several classes of collagens are restricted to metazoans in our data set: fibrillar collagen, a major component of metazoan connective tissues; and collagen IV *sensu stricto* (i.e. excluding MvCN, which has many fewer triple helix repeats than metazoan collagen IV), the major component of metazoan basement membranes. The C-terminal domains of these collagens allow them to form highly oligomeric, cross-linked superstructures—fibrils and networks, respectively—that are key to their roles as the structural scaffold of metazoan ECM (Ricard-Blum, 2011). In addition to proline hydroxylation, metazoan collagens also undergo diverse, functionally important post-translational modifications, such as lysine hydroxylation and cross-linking (Eyre and Wu, 2005; Fidler et al., 2014; Rodriguez-Pascual and Slatter, 2016), proteolytic cleavage (Canty and Kadler, 2005; Ricard-Blum, 2011), and glycosylation (Hennet, 2019), and it remains to be seen whether non-metazoan collagens might undergo similar modifications. Thus, while collagen domains are not unique to Metazoa, it may be the case that innovations related to collagen, such as the origin of fibrillar collagens/collagen IV and their associated superstructures, increased modification with 4-hydroxyproline, and/or innovations in collagen-interacting and collagen-modifying proteins not addressed by our data set, facilitated the evolution of complex multicellularity during metazoan origins.

The ubiquity of collagen domains in choanoflagellates raises the question of what role collagens might be playing in single-celled and colonial relatives of metazoans. In a possible parallel to fibrillar collagen and collagen IV in metazoans, we detected an ancient collagenous lectin, ChCL, that is widespread across choanoflagellates and was likely present in the last common ancestor of all choanoflagellates. Choanoflagellates secrete extracellular matrices that are dynamically regulated across multiple life history stages and play functional roles in substrate adhesion (Dayel and King, 2014; Leadbeater, 2008), mating regulation (Woznica et al., 2017), and multicellular colony formation (Dayel et al., 2011; Larson et al., 2020; Levin et al., 2014). Furthermore, choanoflagellates interact closely with bacteria as predators (Dayel and King, 2014), as signaling partners (Alegado et al., 2012; Ireland et al., 2020; Woznica et al., 2017, 2016), and possibly as symbionts (Hake et al., 2021). We found that some choanoflagellates encode collagens with domains characteristic of metazoan defense collagens, such as scavenger receptors and ficolins, and that the choanoflagellate collagen ChCL contains a domain similar to putative bacterial cell wall proteins, suggestive of a potential role for these collagens in mediating choanoflagellate-bacteria interactions. In general, while non-ECM collagens—i.e., the defense collagens (Casals et al., 2019; Fraser and Tenner, 2008), transmembrane collagens (Franzke et al., 2005; Loria et al., 2004; Maertens et al., 2007), and soluble collagenous signaling proteins (Leclère et al., 2020; Mikkola and Thesleff, 2003; Pyagay et al., 2005)—are relatively well characterized in vertebrates and in some invertebrate model organisms, little is known about non-ECM collagens in basal metazoans. This leaves a gap in our understanding of any ancient non-ECM collagens that may have been present in early metazoan evolution. Future work to characterize the function and localization of collagens in basal metazoans and close metazoan relatives, including choanoflagellates, may shed light on the early roles of collagens both within and outside of the ECM.

## Methods

### Identification of collagen domains and collagen-associated domains in non-metazoan holozoans

To catalogue putative prolyl 4-hydroxylase and collagen domains in holozoans, we obtained the predicted proteomes of 22 choanoflagellates, four filastereans, two ichthyosporeans, and one corallochytrid (Table S1). We performed InterProScan annotation of each proteome using the AgBase InterProScan Singularity container (running InterProScan version 5.41-78.0; available at https://hub.docker.com/r/agbase/interproscan) on the University of California Berkeley Savio computational cluster to obtain Gene3D, Panther, Pfam, Phobius, PrositePatterns, SMART, and SuperFamily annotations of every protein. The Phobius data were used for transmembrane region predictions and signal sequence predictions. Any protein region annotated as IPR008160 (“Collagen triple helix repeat”; https://www.ebi.ac.uk/interpro/entry/InterPro/IPR008160/) was considered a collagen domain, and any region annotated as IPR006620 (“Prolyl 4-hydroxylase, alpha subunit”; https://www.ebi.ac.uk/interpro/entry/InterPro/IPR006620/) was considered a putative prolyl 4-hydroxylase domain. Domains annotated as IPR001442 (“Collagen IV, non-collagenous”; https://www.ebi.ac.uk/interpro/entry/InterPro/IPR001442/) or IPR036954 (“Collagen IV, non-collagenous domain superfamily”; https://www.ebi.ac.uk/interpro/entry/InterPro/IPR036954/) were considered NC1 domains, domains annotated as IPR000885 (“Fibrillar collagen, C-terminal”; https://www.ebi.ac.uk/interpro/entry/InterPro/IPR000885/) were considered COLF1 domains.

### Identification of collagen domains and prolyl 4-hydroxylases in metazoans and non-holozoan eukaryotes

To characterize the distribution of putative prolyl 4-hydroxylase domains and collagen domains across non-holozoan eukaryotes and metazoans, we took advantage of the publicly available proteomes in the UniProtKB database. We first assembled a list of all eukaryotic proteomes available in the UniProtKB database as of June 30th, 2021 (UniProt release 2021_03; https://www.uniprot.org/) by searching the UniProtKB Proteomes database (https://www.uniprot.org/proteomes/) with the query and downloading the results. We filtered out low-quality proteomes using the following criteria: first, we discarded any proteomes with a protein count less than 200; second, we discarded proteomes classified by UniProt’s Complete Proteome Detector (CPD) as “Outlier (low value)”; and third, we discarded proteomes with BUSCO scores lower than 50% (or, for proteomes with no BUSCO score provided, we discarded those with a protein count less than 20,000). These criteria were chosen in order to minimize inclusion of low-quality proteomes while retaining at least one proteome from each major eukaryotic group.

After filtering the proteomes, the “Taxonomic lineage” provided by UniProt for each proteome was then used as the basis for the topologies of all phylogenetic trees presented in this paper, including assignment of each organism to a taxonomic family. At this step, the following modifications were made: (1) We rearranged the base of the eukaryotic phylogeny to match the most up-to-date eukaryotic tree of life based on Burki et al. (2020), and the base of the metazoan phylogeny to reflect known relationships between phyla. (2) We discarded proteomes from organisms labeled “fungal sp. no. X” (where X is a species number) due to unclear phylogenetic placement. (3) For 40 species whose UniProt taxonomic lineage did not include a family name, a family name and taxonomic lineage were assigned from the Global Biodiversity Information Facility (GBIF) database (https://www.gbif.org/); in the 11 out of 40 cases where the GBIF database did not list a family name due to phylogenetic uncertainty, the organism and its proteome was discarded.

Having assembled our eukaryotic phylogeny and list of proteomes for analysis, we next searched for collagen domains in metazoans and non-holozoan eukaryotes by searching the UniProtKB protein database (https://www.uniprot.org/uniprot/) with the following queries:

1. NOT taxonomy:“Opisthokonta [33154]” IPR008160 taxonomy:“Eukaryota [2759]”
2. taxonomy:“Fungi [4751]” IPR008160
3. taxonomy:“Metazoa [33208]” IPR008160

We downloaded all hits for these three queries, then discarded any hits whose UniProt Organism ID was not present in our list of filtered proteomes. To catalogue putative prolyl 4-hydroxylase domains in metazoans and non-holozoan eukaryotes, we repeated the above procedure replacing “IPR008160” with “IPR006620”.

### Quantification of collagen repeat length and proline content

To compare the characteristics of collagens across eukaryotic diversity, we analyzed the collagen repeat length and proline content of collagens in different eukaryotic clades. Collagen repeat length was assessed by quantifying, for each protein, the total number of triplet repeats matching the pattern (Gly-X-Y)_*n*_, where *n* ≥ 2 and X and Y can be any amino acid. The resulting value was either expressed in units of repeats or multiplied by 3 when expressed in units of amino acids. Proline content was assessed by dividing the total number of prolines found in the X and Y positions of those triplet repeats by twice the total number of repeats. For example, a protein with the sequence MGPPGNNGPNNNNPGNPNNNNGPNGPN* has a “collagen repeat length” of 5 repeats or 15 amino acids, and its “proportion of proline in X and Y positions” is 50%.

### Analysis of MvCN domains

We investigated the non-collagenous domains of *Ministeria vibrans* MvCN through a combination of BLAST search and amino acid alignment. First, we conducted a search of the NCBI BLAST database (https://blast.ncbi.nlm.nih.gov/Blast.cgi) using the MvCN N-terminal non-collagenous domain (amino acids 1-731) as a query. This resulted in three hits, NBV83445.1, NTW51976.1, and WP_161718411.1, all of which are bacterial proteins. Alignment of the MvCN N-terminal domain against these hits (Fig. S3) was performed using the Geneious alignment algorithm in Geneious Prime 2021.1.1 (https://www.geneious.com). We aligned the MvCN NC1 domain (amino acids 938-1122) against the NC1 domains of collagen IV from the ctenophore *Mnemiopsis leidyi* (protein ML18198a-PA) and the sponge *Oscarella carmela* (protein Ocar_m.306941) using the same alignment procedure (Fig. S3).

### Analysis of ChCL domains

To investigate the origins of the conserved choanoflagellate collagen ChCL, we first performed a search of the NCBI BLAST database using the entire amino acid sequence of the representative *Salpingoeca urceolata* ChCL protein (Surc_m.480486) as a query. This yielded as hits many metazoan proteins that aligned to the middle non-collagenous domain of the protein (amino acids ∼200-900). We aligned *S. urceolata* ChCL to the top hit, XP_013383323.1, as described above (Fig. S4). We then performed a second search with only the C-terminal domain of *S. urceolata* ChCL (amino acids 920-1136) as a query, which resulted in predominantly bacterial hits, and we aligned the *S. urceolata* ChCL C-terminal domain to the top hit, MSX41286.1, as described above (Fig. S4).

## Supporting information

Supplementary Data Key

## Acknowledgements

We thank O. Muellerklein of Berkeley Research Computing and F. McCarthy and S. Saha of the University of Arizona for advice on InterProScan implementation. This research used the Savio computational cluster resource provided by the Berkeley Research Computing program at the University of California, Berkeley (supported by the UC Berkeley Chancellor, Vice Chancellor for Research, and Chief Information Officer). T.A.L. was supported by an NSF GRFP Fellowship and the Berkeley Fellowship for Graduate Study.

## Notes

### Competing Interest Statement

The authors have declared no competing interest.

