## Supplementary Materials for "Widespread distribution of collagens and collagen-associated domains in eukaryotes"

This PDF file includes:

Figs. S1 to S7...........................................................2
Table S1..................................................................18

Other Supplementary Materials for this manuscript include the following:

SupplementaryDataFile1.csv
SupplementaryDataFile2.csv
SupplementaryDataFile3.cv
SupplementaryDataKey.pdf

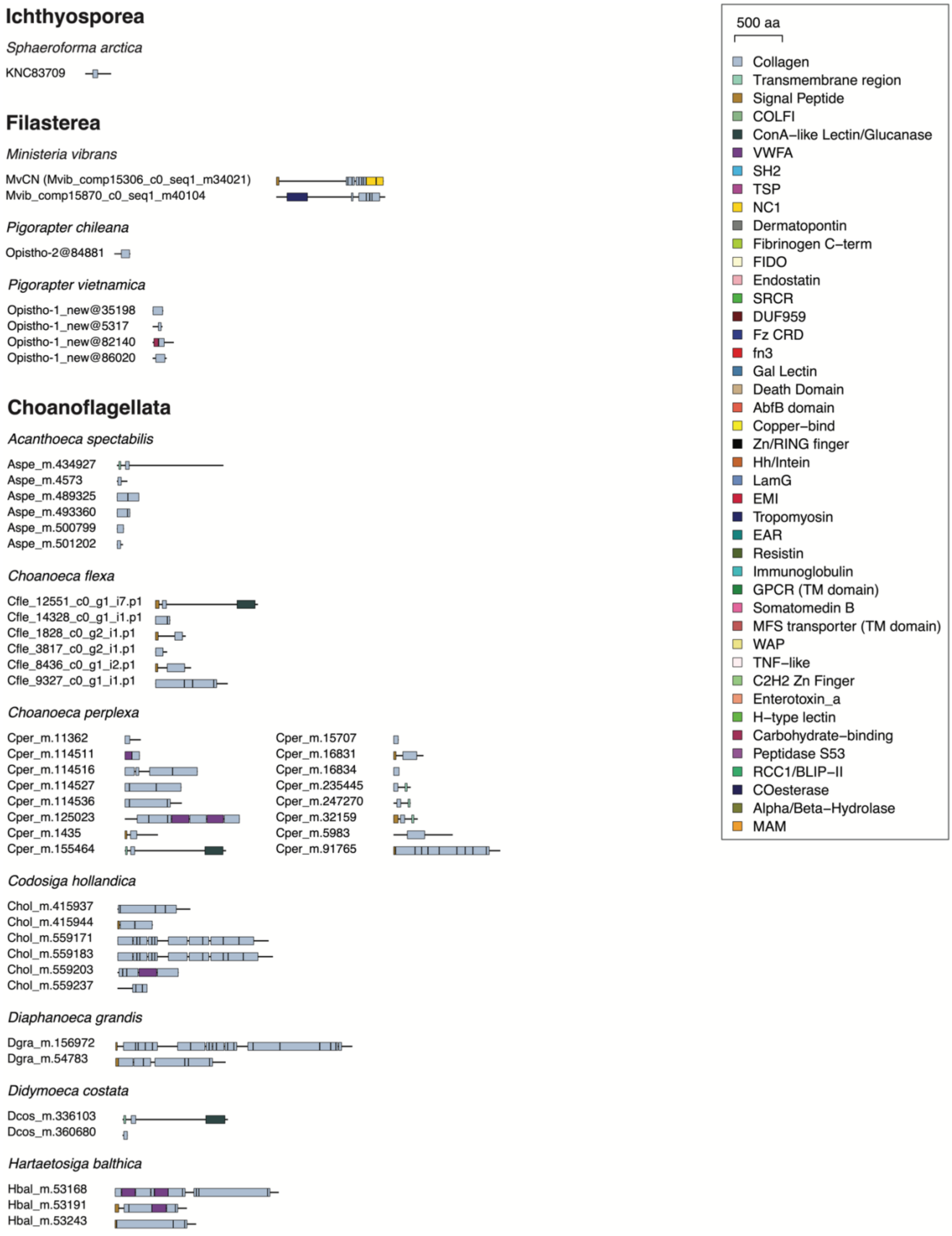
Ichthyosporea
Sphaeroforma arctica
KNC83709
Filasterea
Ministeria vibrans
MvCN (Mvib_comp15306_c0_seq1_m34021)
Mvib_comp15870_c0_seq1_m40104
Pigorapter chileana
Opistho-2@84881
Pigorapter vietnamica
Opistho-1_new@35198
Opistho-1_new@5317
Opistho-1_new@82140
Opistho-1_new@86020
Choanoflagellata
Acanthoeca spectabilis
Aspe_m.434927
Aspe_m.4573
Aspe_m.489325
Aspe_m.493360
Aspe_m.500799
Aspe_m.501202
Choanoeca flexa
Cfle_12551_c0_g1_i7.p1
Cfle_14328_c0_g1_i1.p1
Cfle_1828_c0_g2_i1.p1
Cfle_3817_c0_g2_i1.p1
Cfle_8436_c0_g1_i2.p1
Cfle_9327_c0_g1_i1.p1
Choanoeca perplexa
Cper_m.11362
Cper_m.114511
Cper_m.114516
Cper_m.114527
Cper_m.114536
Cper_m.125023
Cper_m.1435
Cper_m.155464
Cper_m.15707
Cper_m.16831
Cper_m.16834
Cper_m.235445
Cper_m.247270
Cper_m.32159
Cper_m.5983
Cper_m.91765
Codosiga hollandica
Chol_m.415937
Chol_m.415944
Chol_m.559171
Chol_m.559183
Chol_m.559203
Chol_m.559237
Diaphanoeca grandis
Dgra_m.156972
Dgra_m.54783
Didymoeca costata
Dcos_m.336103
Dcos_m.360680
Hartaetosiga balthica
Hbal_m.53168
Hbal_m.53191
Hbal_m.53243
500 aa
Collagen
Transmembrane region
Signal Peptide
COLFI
ConA–like Lectin/Glucanase
VWFA
SH2
TSP
NC1
Dermatopontin
Fibrinogen C–term
FIDO
Endostatin
SRCR
DUF959
Fz CRD
fn3
Gal Lectin
Death Domain
AbfB domain
Copper–bind
Zn/RING finger
Hh/Intein
LamG
EMI
Tropomyosin
EAR
Resistin
Immunoglobulin
GPCR (TM domain)
Somatomedin B
MFS transporter (TM domain)
WAP
TNF–like
C2H2 Zn Finger
Enterotoxin_a
H–type lectin
Carbohydrate–binding
Peptidase S53
RCC1/BLIP–II
COesterase
Alpha/Beta–Hydrolase
MAM

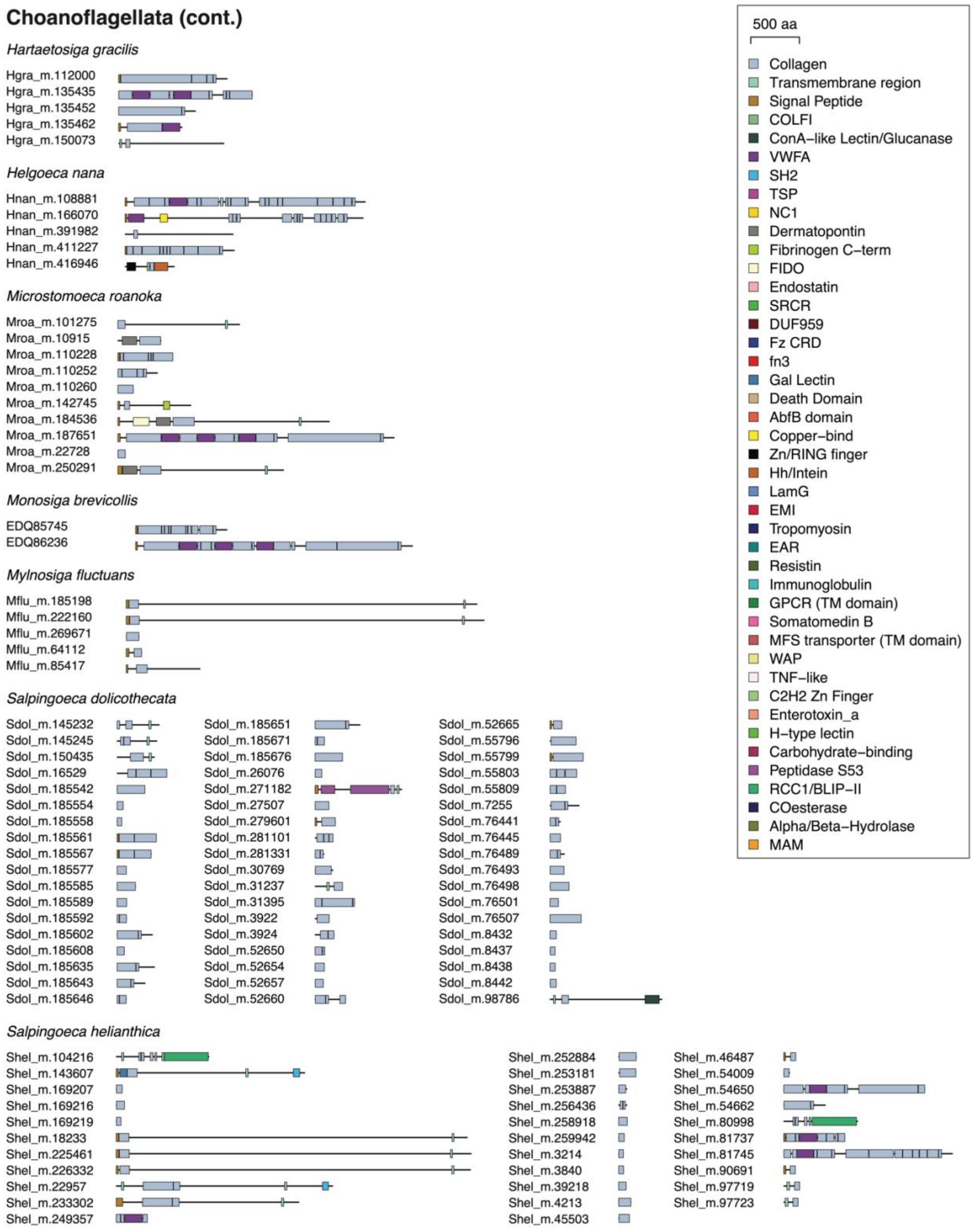
Choanoflagellata (cont.)
Hartaetosiga gracilis
Hgra_m.112000
Hgra_m.135435
Hgra_m.135452
Hgra_m.135462
Hgra_m.150073
Helgoeca nana
Hnan_m.108881
Hnan_m.166070
Hnan_m.391982
Hnan_m.411227
Hnan_m.416946
Microstomoeca roanoka
Mroa_m.101275
Mroa_m.10915
Mroa_m.110228
Mroa_m.110252
Mroa_m.110260
Mroa_m.142745
Mroa_m.184536
Mroa_m.187651
Mroa_m.22728
Mroa_m.250291
Monosiga brevicollis
EDQ85745
EDQ86236
Mylnosiga fluctuans
Mflu_m.185198
Mflu_m.222160
Mflu_m.269671
Mflu_m.64112
Mflu_m.85417
Salpingoeca dolichothecata
Sdol_m.145232
Sdol_m.145245
Sdol_m.150435
Sdol_m.16529
Sdol_m.185542
Sdol_m.185554
Sdol_m.185558
Sdol_m.185561
Sdol_m.185567
Sdol_m.185577
Sdol_m.185585
Sdol_m.185589
Sdol_m.185592
Sdol_m.185602
Sdol_m.185608
Sdol_m.185635
Sdol_m.185643
Sdol_m.185646
Sdol_m.185651
Sdol_m.185671
Sdol_m.185676
Sdol_m.26076
Sdol_m.271182
Sdol_m.27507
Sdol_m.279601
Sdol_m.281101
Sdol_m.281331
Sdol_m.30769
Sdol_m.31237
Sdol_m.31395
Sdol_m.3922
Sdol_m.3924
Sdol_m.52650
Sdol_m.52654
Sdol_m.52657
Sdol_m.52660
Sdol_m.52665
Sdol_m.55796
Sdol_m.55799
Sdol_m.55803
Sdol_m.55809
Sdol_m.7255
Sdol_m.76441
Sdol_m.76445
Sdol_m.76489
Sdol_m.76493
Sdol_m.76498
Sdol_m.76501
Sdol_m.76507
Sdol_m.8432
Sdol_m.8437
Sdol_m.8438
Sdol_m.8442
Sdol_m.98786
Salpingoeca helianthica
Shel_m.104216
Shel_m.143607
Shel_m.169207
Shel_m.169216
Shel_m.169219
Shel_m.18233
Shel_m.225461
Shel_m.226332
Shel_m.22957
Shel_m.233302
Shel_m.249357
Shel_m.252884
Shel_m.253181
Shel_m.253887
Shel_m.256436
Shel_m.258918
Shel_m.259942
Shel_m.3214
Shel_m.3840
Shel_m.39218
Shel_m.4213
Shel_m.45503
Shel_m.46487
Shel_m.54009
Shel_m.54650
Shel_m.54662
Shel_m.80998
Shel_m.81737
Shel_m.81745
Shel_m.90691
Shel_m.97719
Shel_m.97723
500 aa
Collagen
Transmembrane region
Signal Peptide
COLFI
ConA–like Lectin/Glucanase
VWFA
SH2
TSP
NC1
Dermatopontin
Fibrinogen C–term
FIDO
Endostatin
SRCR
DUF959
Fz CRD
fn3
Gal Lectin
Death Domain
AbfB domain
Copper–bind
Zn/RING finger
Hh/Intein
LamG
EMI
Tropomyosin
EAR
Resistin
Immunoglobulin
GPCR (TM domain)
Somatomedin B
MFS transporter (TM domain)
WAP
TNF–like
C2H2 Zn Finger
Enterotoxin_a
H–type lectin
Carbohydrate–binding
Peptidase S53
RCC1/BLIP–II
COesterase
Alpha/Beta–Hydrolase
MAM

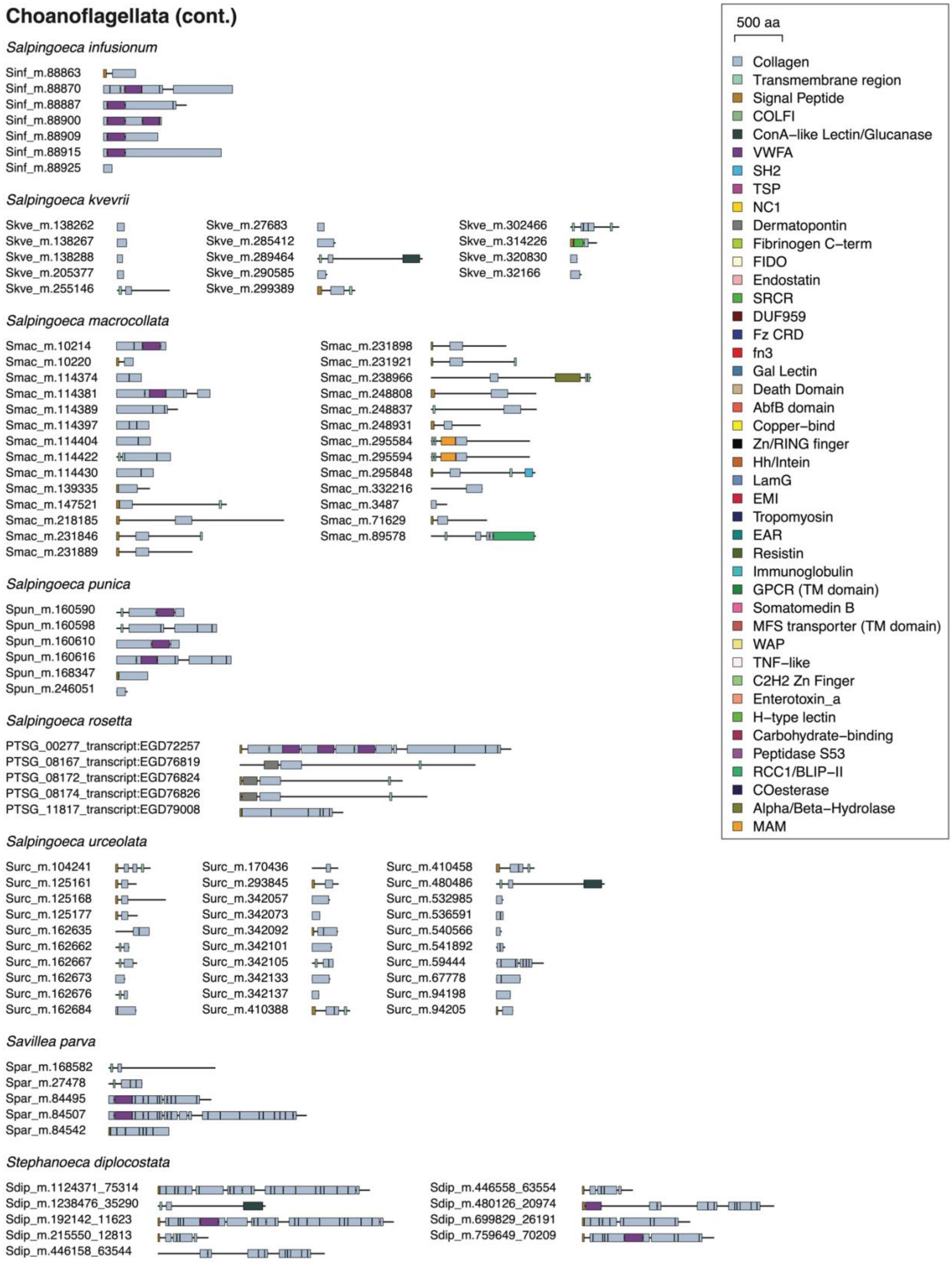
Choanoflagellata (cont.)
Salpingoeca infusionum
Sinf_m.88863
Sinf_m.88870
Sinf_m.88887
Sinf_m.88900
Sinf_m.88909
Sinf_m.88915
Sinf_m.88925
Salpingoeca kvevrii
Skve_m.138262
Skve_m.138267
Skve_m.138288
Skve_m.205377
Skve_m.255146
Skve_m.27683
Skve_m.285412
Skve_m.289464
Skve_m.290585
Skve_m.299389
Skve_m.302466
Skve_m.314226
Skve_m.320830
Skve_m.32166
Salpingoeca macrocollata
Smac_m.10214
Smac_m.10220
Smac_m.114374
Smac_m.114381
Smac_m.114389
Smac_m.114397
Smac_m.114404
Smac_m.114422
Smac_m.114430
Smac_m.139335
Smac_m.147521
Smac_m.218185
Smac_m.231846
Smac_m.231889
Smac_m.231898
Smac_m.231921
Smac_m.238966
Smac_m.248808
Smac_m.248837
Smac_m.248931
Smac_m.295584
Smac_m.295594
Smac_m.295848
Smac_m.332216
Smac_m.3487
Smac_m.71629
Smac_m.89578
Salpingoeca punica
Spun_m.160590
Spun_m.160598
Spun_m.160610
Spun_m.160616
Spun_m.168347
Spun_m.246051
Salpingoeca rosetta
PTSG_00277_transcript:EGD72257
PTSG_08167_transcript:EGD76819
PTSG_08172_transcript:EGD76824
PTSG_08174_transcript:EGD76826
PTSG_11817_transcript:EGD79008
Salpingoeca urceolata
Surc_m.104241
Surc_m.125161
Surc_m.125168
Surc_m.125177
Surc_m.162635
Surc_m.162662
Surc_m.162667
Surc_m.162673
Surc_m.162676
Surc_m.162684
Surc_m.170436
Surc_m.293845
Surc_m.342057
Surc_m.342073
Surc_m.342092
Surc_m.342101
Surc_m.342105
Surc_m.342133
Surc_m.342137
Surc_m.410388
Surc_m.410458
Surc_m.480486
Surc_m.532985
Surc_m.536591
Surc_m.540566
Surc_m.541892
Surc_m.59444
Surc_m.67778
Surc_m.94198
Surc_m.94205
Savillea parva
Spar_m.168582
Spar_m.27478
Spar_m.84495
Spar_m.84507
Spar_m.84542
Stephanoeca diplocostata
Sdip_m.1124371_75314
Sdip_m.1238476_35290
Sdip_m.192142_11623
Sdip_m.215550_12813
Sdip_m.446158_63544
Sdip_m.446558_63554
Sdip_m.480126_20974
Sdip_m.699829_26191
Sdip_m.759649_70209
500 aa
Collagen
Transmembrane region
Signal Peptide
COLFI
ConA–like Lectin/Glucanase
VWFA
SH2
TSP
NC1
Dermatopontin
Fibrinogen C–term
FIDO
Endostatin
SRCR
DUF959
Fz CRD
fn3
Gal Lectin
Death Domain
AbfB domain
Copper–bind
Zn/RING finger
Hh/Intein
LamG
EMI
Tropomyosin
EAR
Resistin
Immunoglobulin
GPCR (TM domain)
Somatomedin B
MFS transporter (TM domain)
WAP
TNF–like
C2H2 Zn Finger
Enterotoxin_a
H–type lectin
Carbohydrate–binding
Peptidase S53
RCC1/BLIP–II
COesterase
Alpha/Beta–Hydrolase
MAM

**Figure S1. Collagen repertoires of 29 non-metazoan holozoans.** Schematic representations of all collagens proteins detected in 22 choanoflagellates, 4 filastereans, 2 ichthyosporeans, and 1 corallochytrid using InterProScan annotation. Protein diagrams were generated in R using the make_prot_fig script (Linden, 2021; https://github.com/tesslinden/interpro-scripts).

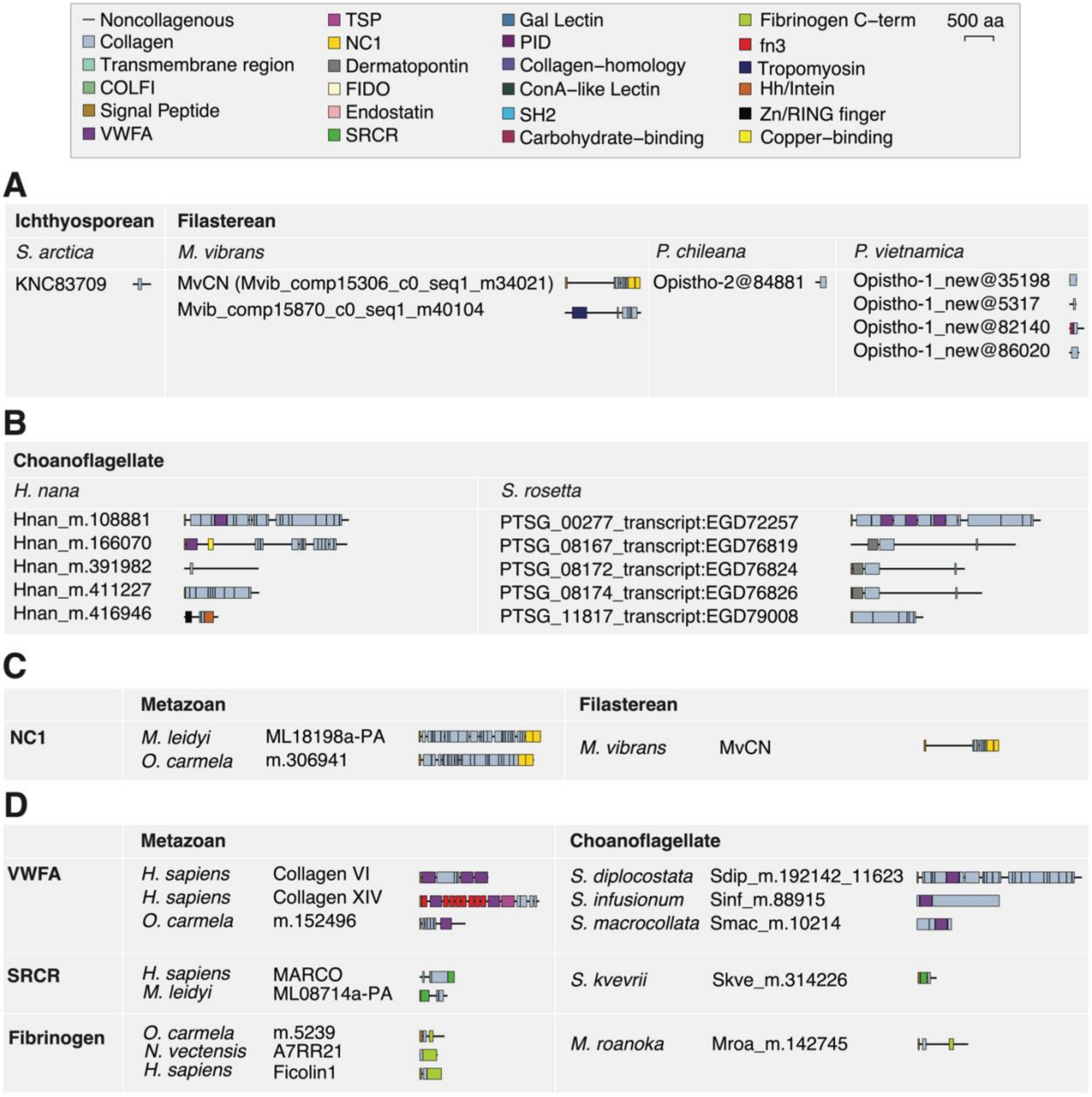

Noncollagenous
Collagen
Transmembrane region
COLFI
Signal Peptide
VWFA
TSP
NC1
Dermatopontin
FIDO
Endostatin
SRCR
Gal Lectin
PID
Collagen–homology
ConA–like Lectin
SH2
Carbohydrate–binding
Fibrinogen C–term
fn3
Tropomyosin
Hh/Intein
Zn/RING finger
Copper–binding
500 aa
A
Ichthyosporean
Filasterean
S. arctica
KNC83709
M. vibrans
MvCN (Mvib_comp15306_c0_seq1_m34021)
Mvib_comp15870_c0_seq1_m40104
P. chileana
Opistho-2@84881
P. vietnamica
Opistho-1_new@35198
Opistho-1_new@5317
Opistho-1_new@82140
Opistho-1_new@86020
B
Choanoflagellate
H. nana
Hnan_m.108881
Hnan_m.166070
Hnan_m.391982
Hnan_m.411227
Hnan_m.416946
S. rosetta
PTSG_00277_transcript:EGD72257
PTSG_08167_transcript:EGD76819
PTSG_08172_transcript:EGD76824
PTSG_08174_transcript:EGD76826
PTSG_11817_transcript:EGD79008
C
Metazoan
Filasterean
NC1
M. leidyi
ML18198a-PA
O. carmela
m.306941
M. vibrans
MvCN
D
Metazoan
Choanoflagellate
VWFA
H. sapiens
Collagen VI
H. sapiens
Collagen XIV
O. carmela
m.152496
S. diplocostata
Sdip_m.192142_11623
S. infusionum
Sinf_m.88915
S. macrocollata
Smac_m.10214
SRCR
H. sapiens
MARCO
M. leidyi
ML08714a-PA
S. kvevrii
Skve_m.314226
Fibrinogen
O. carmela
m.5239
N. vectensis
A7RR21
H. sapiens
Ficolin1
M. roanoka
Mroa_m.142745

**Figure S2. Some holozoan collagens contain known metazoan collagen-associated domains.** **(A)** Schematic representations of the complete collagen repertoires of the ichthyosporean *Sphaeroforma arctica* and the filastereans *Ministeria vibrans, Pigoraptor chileana,* and *Pigoraptor vietnamica.* **(B)** Schematic representations of the complete collagen repertoires of the representative loricate choanoflagellate *Helgoeca nana* and the representative craspedid choanoflagellate *Salpingoeca rosetta*. **(C)** The filasterean *Ministeria vibrans* encodes a protein (MvCN) with collagen domains and an NC1 domain, shown here next to two examples of metazoan collagen IV. **(D)** Many choanoflagellates encode collagens with VWFA domains, a common collagen-associated domain in metazoans. Note that not all choanoflagellate VWFA domain-containing collagens are shown here. The choanoflagellate *Salpingoeca kvevrii* encodes a collagen with an SRCR domain, similar to metazoan scavenger receptors. The choanoflagellate *Microstomoeca roanoka* encodes a collagen with a Fibrinogen domain, similar to metazoan ficolins. *Mnemiopsis leidyi* protein sequences were obtained from Ensembl Metazoa (https://metazoa.ensembl.org/Mnemiopsis_leidyi/). *Oscarella carmela* protein sequences were obtained from Compagen (http://www.compagen.org/datasets.html). *Homo sapiens* and *Nematostella vectensis* protein sequences were obtained from UniProtKB (https://www.uniprot.org/uniprot/). Protein diagrams were generated in R using the make_prot_fig script (Linden, 2021; https://github.com/tesslinden/interpro-scripts).

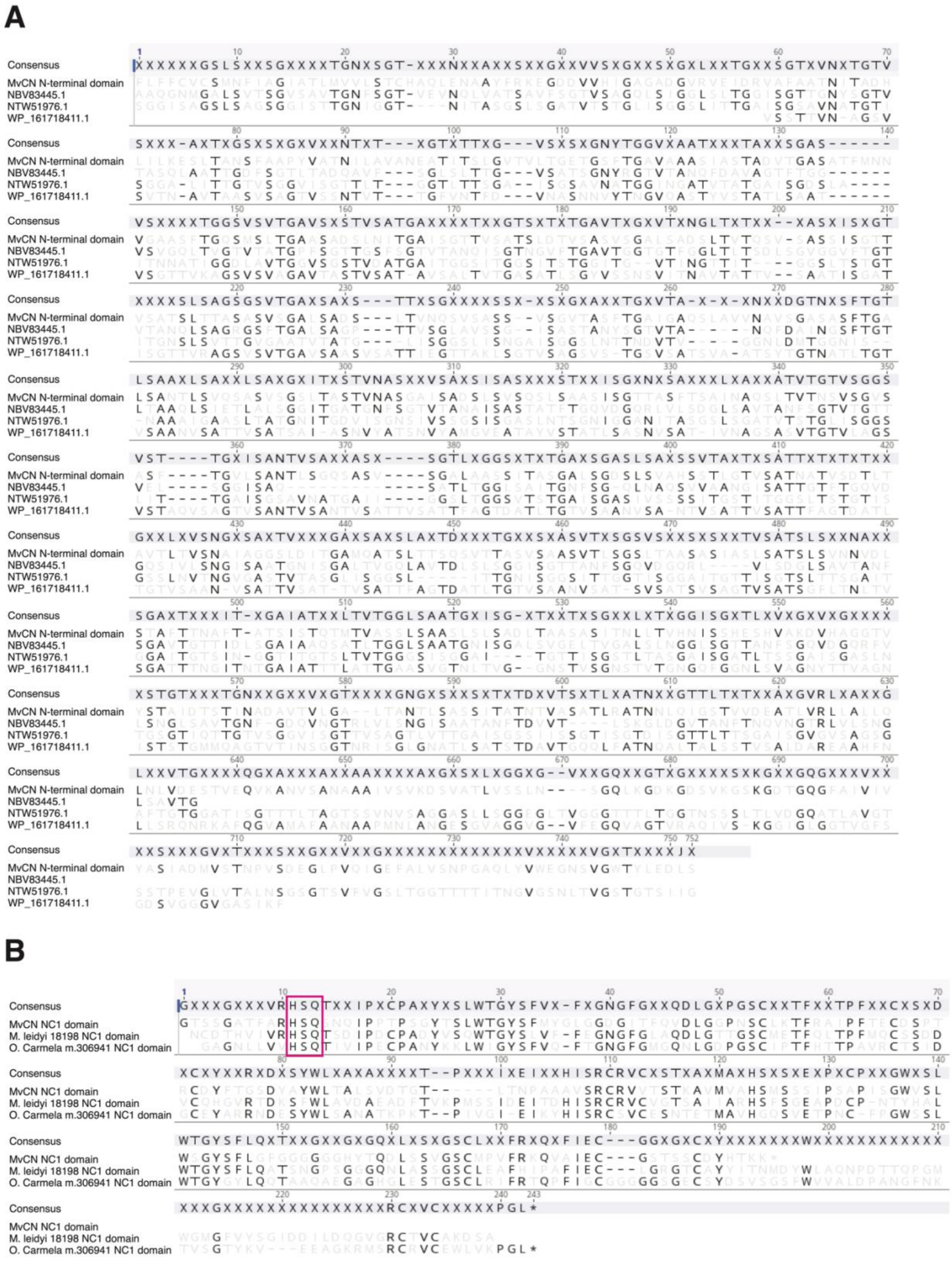

**Figure S3. The *Ministeria vibrans* collagen MvCN is a bacterial fusion protein. (A)** Alignment of the N-terminal non-collagenous domain of MvCN (amino acids 1-731) against its top three BLAST hits, all of which are bacterial proteins. **(B)** Alignment of the MvCN NC1 domain (amino acids 938-1,122) against the NC1 domains of collagen IV in two basal metazoans, the ctenophore *Mnemiopsis leidyi* (protein ML18198a-PA) and the sponge *Oscarella carmela* (protein m.306941). The pink box highlights the conserved HSQ motif that is found in the NC1 domains of collagen IVs but not those of spongins (Fidler et al., 2017). Alignments were performed using the Geneious alignment algorithm in Geneious Prime 2021.1.1 (https://www.geneious.com).

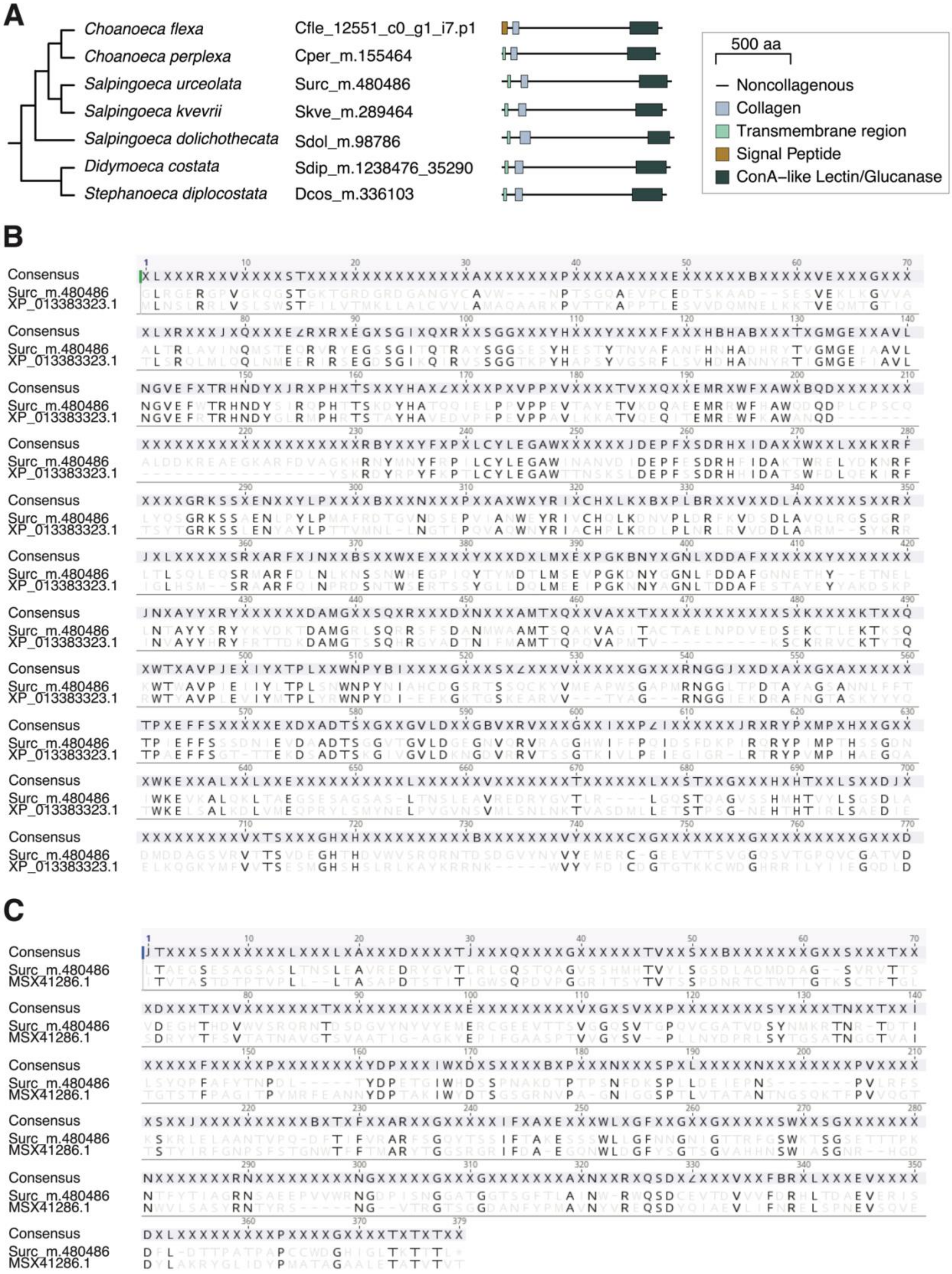
A
Choanoeca flexa
Choanoeca perplexa
Salpingoeca urceolata
Salpingoeca kvevrii
Salpingoeca dolichothecata
Didymoeca costata
Stephanoeca diplocostata
Cfle_12551_c0_g1_i7.p1
Cper_m.155464
Surc_m.480486
Skve_m.289464
Sdol_m.98786
Sdip_m.1238476_35290
Dcos_m.336103
500 aa
Noncollagenous
Collagen
Transmembrane region
Signal Peptide
ConA–like Lectin/Glucanase
B
Consensus
Surc_m.480486
XP_013383323.1
C
Consensus
Surc_m.480486
MSX41286.1

**Figure S4. The choanoflagellate-specific collagen ChCL is a bacterial fusion protein. (A)** Schematic representations of all seven ChCL proteins detected across 22 species of choanoflagellates, most of which are predicted to have an N-terminal transmembrane region, and all of which have an extracellular region consisting of a collagen domain, a non-collagenous domain, and a C-terminal Concanavalin A-like lectin/glucanase domain. These species are broadly distributed across the choanoflagellate phylogeny (see Figure 1), suggesting that ChCL may have been present in the most recent common ancestor of choanoflagellates. Protein diagrams were generated in R using the make_prot_fig script (Linden, 2021; https://github.com/tesslinden/interpro-scripts). **(B)** Alignment of the non-collagenous region (amino acids 144-899) of a representative ChCL from *Salpingoeca urceolata* against its top BLAST hit, a non-collagenous metazoan protein from *Lingula anatina*. **(C)** Alignment of the C-terminal domain (amino acids 776-1,135) of *S. urceolata* ChCL against its top BLAST hit, a protein from an Actinobacterium. Alignments were performed using the Geneious alignment algorithm in Geneious Prime 2021.1.1 (https://www.geneious.com).

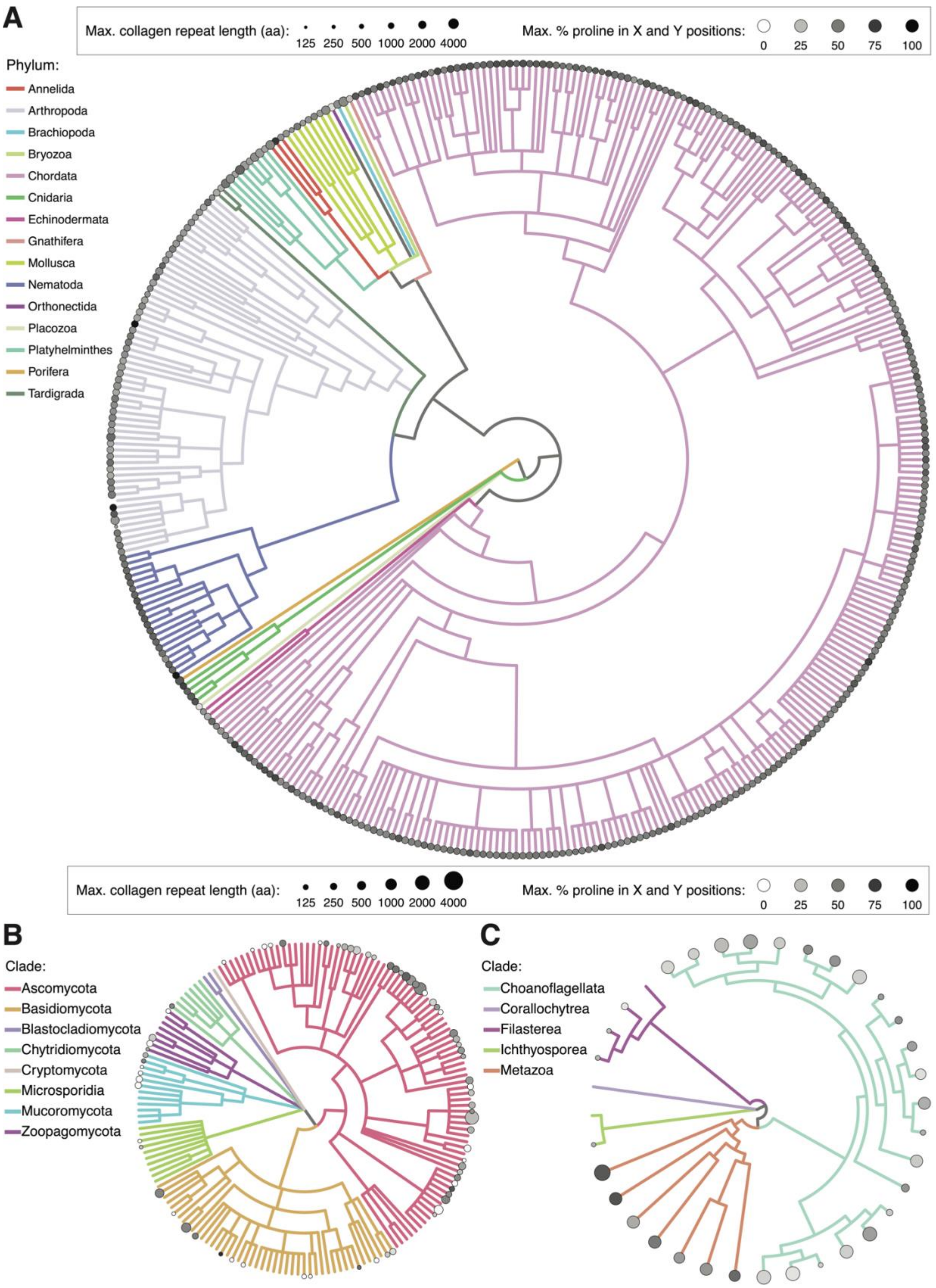
A
Max. collagen repeat length (aa):
125 250 500 1000 2000 4000
Max. % proline in X and Y positions:
0 25 50 75 100
Phylum:
Annelida
Arthropoda
Brachiopoda
Bryozoa
Chordata
Cnidaria
Echinodermata
Gnathifera
Mollusca
Nematoda
Orthonectida
Placozoa
Platyhelminthes
Porifera
Tardigrada
Max. collagen repeat length (aa):
125 250 500 1000 2000 4000
Max. % proline in X and Y positions:
0 25 50 75 100
B
Clade:
Ascomycota
Basidiomycota
Blastocladiomycota
Chytridiomycota
Cryptomycota
Microsporidia
Mucoromycota
Zoopagomycota
C
Clade:
Choanoflagellata
Corallochytrea
Filasterea
Ichthyosporea
Metazoa

**Figure S5. Collagen domains across metazoan, fungal, and holozoan diversity. (A,B)** In these trees, each stem represents one taxonomic family for which at least one proteome meeting our quality criteria (see Methods) is present in the UniProtKB database. The presence of a circle indicates that at least one species in that family encodes a collagen domain (IPR008160). The size of the circle represents the maximum collagen repeat length of any collagen in the family, while the shading of the circle represents the maximum proline content of any collagen in the family (see Methods). Note that the maximum collagen repeat length and maximum proline content may be from different proteins**,** and that the length scale is different between the top and bottom panels, as indicated in the key. **(A)** Collagen domains across metazoan families, color-coded by phylum. **(B)** Collagen domains across fungal families, color-coded by phylum. **(C)** In this tree of holozoan species, each stem represents a single species whose proteome was investigated for collagen domains. The presence of a circle indicates that the species encodes a collagen domain. The size of the circle represents the maximum collagen repeat length of any collagen in the species, while the shading of the circle represents the maximum proline content of any collagen in the species; again, note that the maximum collagen repeat length and maximum proline content may be from different proteins. The seven metazoan species shown here (*Daphnia pulex, Branchiostoma floridae, Nematostella vectensis, Strongylocentrotus purpuratus, Amphimedon queenslandica, Capitella teleta,* and *Lottia gigantea)* were subsetted from our UniProtKB analysis and were selected to represent seven maximally diverse phyla while also maximizing ancestral gene family retention based on the analysis of Richter et al. (2018).

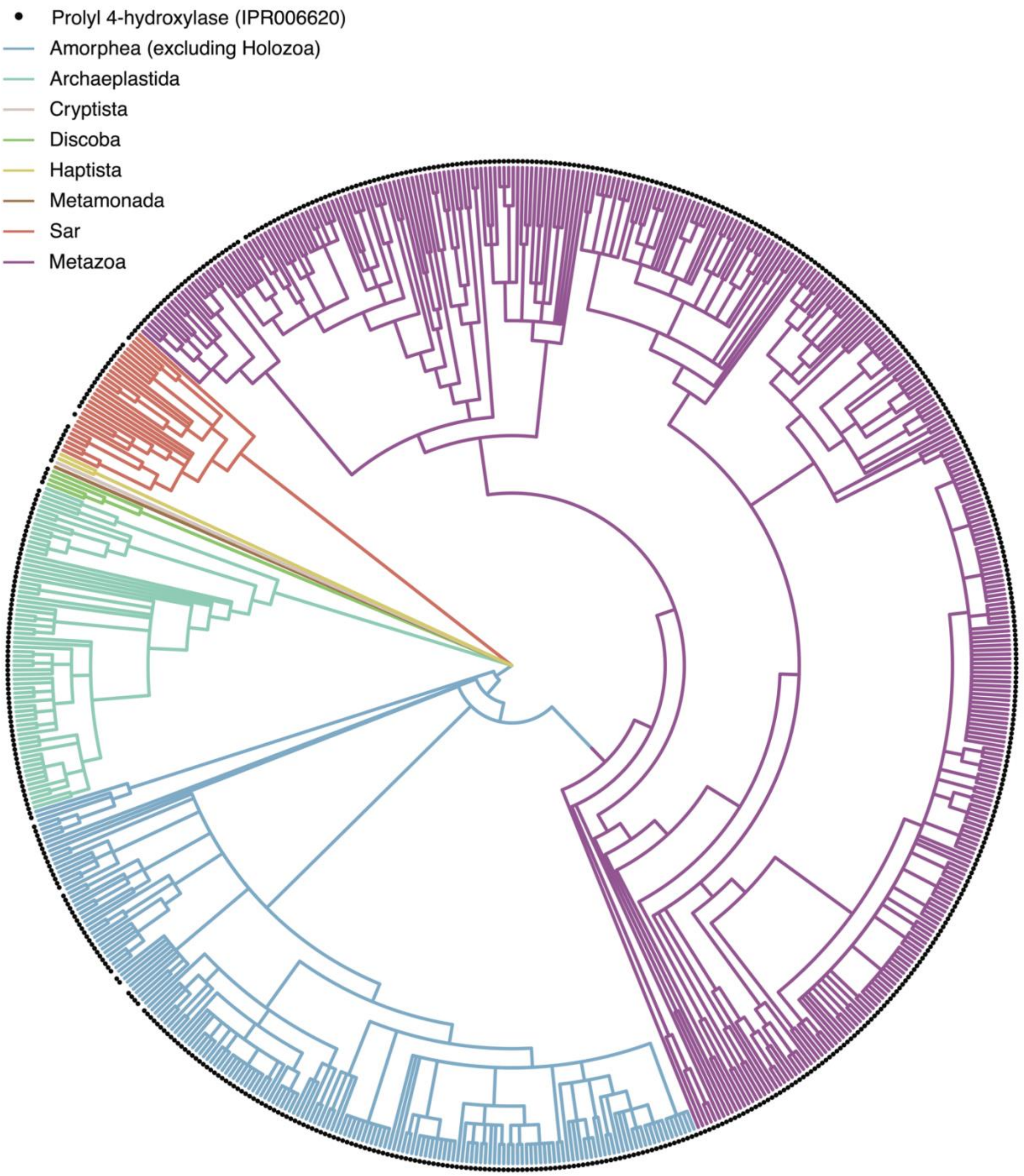
Prolyl 4-hydroxylase (IPR006620)
Amorphea (excluding Holozoa)
Archaeplastida
Cryptista
Discoba
Haptista
Metamonada
Sar
Metazoa

**Figure S6. Distribution of putative prolyl 4-hydroxylase domains across eukaryotes.** We searched for putative prolyl 4-hydroxylase domains (IPR00620) across all eukaryotic proteomes in the UniProtKB database. Each branch of this phylogenetic tree represents a single taxonomic family represented by at least one proteome in UniProtKB that met our quality cutoffs (see Methods). The presence of a black circle indicates that at least one proteome within that family encodes a prolyl 4-hydroxylase domain. The topology of the tree is identical to the trees shown in Fig. 2C-F and Fig. S5.

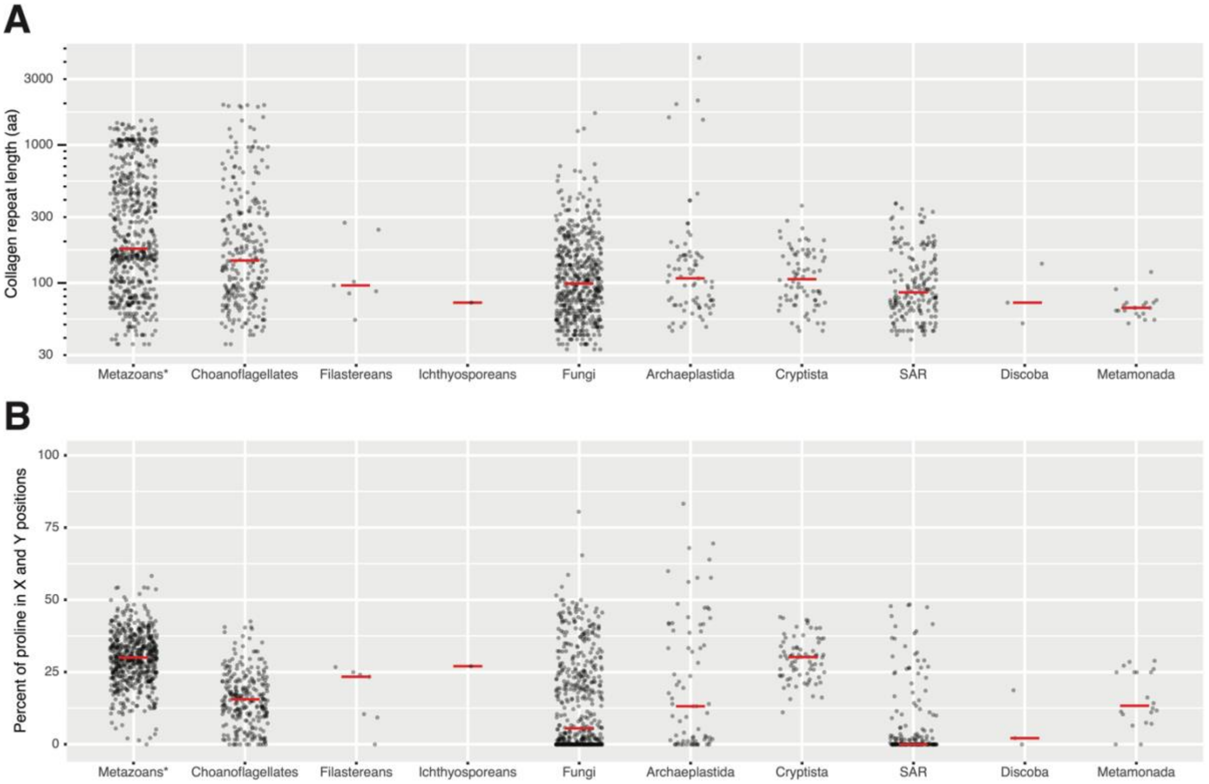
A
Collagen repeat length (aa)
3000
1000
300
100
30
Metazoans*
Choanoflagellates
Filastereans
Ichthyosporeans
Fungi
Archaeplastida
Cryptista
SAR
Discoba
Metamonada
B
Percent of proline in X and Y positions
100
75
50
25
0
Metazoans*
Choanoflagellates
Filastereans
Ichthyosporeans
Fungi
Archaeplastida
Cryptista
SAR
Discoba
Metamonada

**Figure S7. Collagen domain length and proline content across eukaryotic diversity.** We examined **(A)** collagen repeat length and **(B)** proline content (see Methods) across all collagens detected in metazoans (*n* = 57,104 collagens), choanoflagellates (*n* = 258 collagens); filastereans (*n* = 7 collagens); ichthyosporeans (*n* = 1 collagen); fungi (*n* = 508 collagens); Archaeplastida (*n* = 75 collagens); Cryptista (*n* = 76 collagens); SAR (*n* = 176 collagens); Discoba (*n* = 3 collagens); and Metamonada (*n* = 20 collagens). Note that the y-axis for panel A is on a log-scale. Red bars represent medians. *: to more clearly visualize the distribution of the metazoan collagen data, a randomly selected 1% of metazoan collagens are shown here.

Table S1. *Holozoan proteome sources.*

| Species | Basis for proteome prediction‡ | Source |
|---|---|---|
| ***Acanthoeca spectabilis*** | T | (Richter et al., 2018) (https://figshare.com/articles/dataset/Data_from_The_ancestral_animal_genetic_toolkit_revealed_by_diverse_choanoflagellate_Ts/5686984?file=9955288) |
| ***Capsaspora owczarzaki*** | G | Ensembl Protist (https://protists.ensembl.org/Capsaspora_owczarzaki_atcc_30864_gca_000151315) |
| ***Choanoeca flexa*** | T | (Brunet et al., 2019) (https://figshare.com/articles/dataset/Choanoeca_flexa_nonredundant_predicted_proteins_fasta/8216291?file=15312674) |
| ***Choanoeca perplexa*** | T | (Richter et al., 2018) (https://figshare.com/articles/dataset/Data_from_The_ancestral_animal_genetic_toolkit_revealed_by_diverse_choanoflagellate_transcriptomes/5686984?file=9955288) |
| ***Codosiga hollandica*** | T | (Richter et al., 2018) (https://figshare.com/articles/dataset/Data_from_The_ancestral_animal_genetic_toolkit_revealed_by_diverse_choanoflagellate_transcriptomes/5686984?file=9955288) |
| ***Creolimax fragrantissima*** | G | (de Mendoza et al., 2015) (https://figshare.com/articles/dataset/Creolimax_fragrantissima_genome_data/1403592?file=3328016) |
| ***Didymoeca costata*** | T | (Richter et al., 2018) (https://figshare.com/articles/dataset/Data_from_The_ancestral_animal_genetic_toolkit_revealed_by_diverse_choanoflagellate_transcriptomes/5686984?file=9955288) |
| ***Hartaetosiga balthica*** | T | (Richter et al., 2018) (https://figshare.com/articles/dataset/Data_from_The_ancestral_animal_genetic_toolkit_revealed_by_diverse_choanoflagellate_transcriptomes/5686984?file=9955288) |
| ***Hartaetosiga gracilis*** | T | (Richter et al., 2018) (https://figshare.com/articles/dataset/Data_from_The_ancestral_animal_genetic_toolkit_revealed_by_diverse_choanoflagellate_transcriptomes/5686984?file=9955288) |
| ***Helgoeca nana*** | T | (Richter et al., 2018) (https://figshare.com/articles/dataset/Data_from_The_ancestral_animal_genetic_toolkit_revealed_by_divers |

| | | |
|---|---|---|
| | | e_choanoflagellate_transcriptomes/5686984?file=9955288) |
| ***Microstomoeca roanoka*** | T | (Richter et al., 2018) (https://figshare.com/articles/dataset/Data_from_The_ancestral_animal_genetic_toolkit_revealed_by_diverse_choanoflagellate_transcriptomes/5686984?file=9955288) |
| ***Ministeria vibrans*** | T | (Grau-Bové et al., 2017) |
| ***Monosiga brevicollis*** | G | Ensembl Protist (https://protists.ensembl.org/Monosiga_brevicollis_mx1_gca_000002865) |
| ***Mylnosiga fluctuans*** | T | (Richter et al., 2018) (https://figshare.com/articles/dataset/Data_from_The_ancestral_animal_genetic_toolkit_revealed_by_diverse_choanoflagellate_transcriptomes/5686984?file=9955288) |
| ***Pigoraptor chileana*** | T | (Hehenberger et al., 2017) (https://datadryad.org/stash/dataset/doi:10.5061/dryad.26bv4) |
| ***Pigoraptor vietnamica*** | T | (Hehenberger et al., 2017) (https://datadryad.org/stash/dataset/doi:10.5061/dryad.26bv4) |
| ***Salpingoeca dolichothecata*** | T | (Richter et al., 2018) (https://figshare.com/articles/dataset/Data_from_The_ancestral_animal_genetic_toolkit_revealed_by_diverse_choanoflagellate_transcriptomes/5686984?file=9955288) |
| ***Salpingoeca helianthica*** | T | (Richter et al., 2018) (https://figshare.com/articles/dataset/Data_from_The_ancestral_animal_genetic_toolkit_revealed_by_diverse_choanoflagellate_transcriptomes/5686984?file=9955288) |
| ***Salpingoeca infusionum*** | T | (Richter et al., 2018) (https://figshare.com/articles/dataset/Data_from_The_ancestral_animal_genetic_toolkit_revealed_by_diverse_choanoflagellate_transcriptomes/5686984?file=9955288) |
| ***Salpingoeca kvevrii*** | T | (Richter et al., 2018) (https://figshare.com/articles/dataset/Data_from_The_ancestral_animal_genetic_toolkit_revealed_by_diverse_choanoflagellate_transcriptomes/5686984?file=9955288) |
| ***Salpingoeca macrocollata*** | T | (Richter et al., 2018) (https://figshare.com/articles/dataset/Data_from_The_ancestral_animal_genetic_toolkit_revealed_by_divers |

| | | |
|---|---|---|
| | | e_choanoflagellate_transcriptomes/5686984?file=9955288) |
| ***Salpingoeca punica*** | T | (Richter et al., 2018) (https://figshare.com/articles/dataset/Data_from_The_ancestral_animal_genetic_toolkit_revealed_by_diverse_choanoflagellate_transcriptomes/5686984?file=9955288) |
| ***Salpingoeca rosetta*** | G | Ensembl Protist (https://protists.ensembl.org/Salpingoeca_rosetta_gca_000188695) |
| ***Salpingoeca urceolata*** | T | (Richter et al., 2018) (https://figshare.com/articles/dataset/Data_from_The_ancestral_animal_genetic_toolkit_revealed_by_diverse_choanoflagellate_transcriptomes/5686984?file=9955288) |
| ***Savillea parva*** | T | (Richter et al., 2018) (https://figshare.com/articles/dataset/Data_from_The_ancestral_animal_genetic_toolkit_revealed_by_diverse_choanoflagellate_transcriptomes/5686984?file=9955288) |
| ***Sphaeroforma arctica*** | G | Ensembl Protist (https://protists.ensembl.org/Sphaeroforma_arctica_jp610_gca_001186125/) |
| ***Stephanoeca diplocostata*** | T | (Richter et al., 2018) (https://figshare.com/articles/dataset/Data_from_The_ancestral_animal_genetic_toolkit_revealed_by_diverse_choanoflagellate_transcriptomes/5686984?file=9955288) (Note: Australian and French proteomes were concatenated into one proteome for this study) |
| ***Syssomonas multiformis*** | T | (Hehenberger et al., 2017) (https://datadryad.org/stash/dataset/doi:10.5061/dryad.26bv4) |

‡T = transcriptome; G = genome
