## Supplementary Data Key for "Widespread distribution of collagens and collagen-associated domains in eukaryotes"

1. SupplementaryDataFile1.csv

Description: Data for all collagen domains detected across all eukaryotes analyzed for this study. Each row represents a single collagen domain, i.e., an uninterrupted stretch of repeats with the pattern (Gly-X-Y)n.

'binom': The Latin name for the species in which this collagen domain was detected. For non-holozoans and metazoans, the name is exactly as found in the UniProt database.
'pathString': A list of all parent taxa leading to this species in the phylogenetic tree found in the all_edges file (for non-holozoans and metazoans) or in the holozoan_edges file (for non-metazoan holozoans).
'level1': The first taxon of the pathString.
'level2': The second taxon of the pathString.
'level3': The third taxon of the pathString.
'level4': The fourth taxon of the pathString.
'level5': The fifth taxon of the pathString.
'level6': The sixth taxon of the pathString.
'level7': The seventh taxon of the pathString.
'family': The family to which this species belongs. NA for non-metazoan holozoans.
'genus': The genus to which this species belongs.
'species': The species name.
'uniprot_organism_id': The UniProt organism ID for the proteome in which this domain was detected. NA for non-metazoan holozoans.
'uniprot_protein_name': The full name of the protein containing this domain as recorded in the UniProt database. NA for non-metazoan holozoans.
'protein_id': A unique protein identifier. For non-holozoans and metazoans, this corresponds to the UniProt protein id.
'start': The amino acid number at which this domain begins.
'stop': The amino acid number at which this domain ends.
'length': The length of this domain, in amino acids (i.e., [stop]-[start]+1).
'protein_length': The length of the protein containing this domain, in amino acids.
'p2': The proportion of prolines in position X in this domain.
'p3': The proportion of prolines in position Y in this domain.
'pavg': The proportion of prolines in positions X and Y in this domain.
'protein_pavg': The total proportion of prolines in positions X and Y across all collagen domains for the protein containing this domain.
'total_cthr_length': The total collagen repeat length of the protein containing this domain (i.e., the sum of the lengths of all collagen domains in this protein).
'max_unint_cthr_length': The maximum length of any single collagen domain in the protein containing this domain.
'fam_max_total_cthr_length': The maximum value of total_cthr_length for any collagen detected in this organism's family. NA for non-metazoan holozoans.
'fam_max_pavg': The maximum value of protein_pavg for any collagen detected in this organism's family. NA for non-metazoan holozoans.

'has_ph': Was a putative prolyl 4-hydroxylase domain detected in the proteome(s) of the organism that encodes this collagen domain? TRUE = yes; FALSE = no.

2. SupplementaryDataFile2.csv

Description: Data for the phylogenetic trees represented in Fig. 1 and Fig. S5C. Each row represents a node of the phylogenetic tree. Nodes without an official name have been arbitrarily assigned a letter of the alphabet as a unique identifier.

'from': The parent taxon of the node that this row represents.
'to': The name of the node that this row represents. For metazoan leaf nodes, the name is exactly as found in the UniProt database.
'holozoa_clade': The holozoan clade to which this node belongs: Metazoa, Choanoflagellata, Filasterea, Corallochytrea, or Ichthyosporea. Nodes that are supertaxa of these clades receive the value 'Roots'.
'has_ph': If this node is a leaf node, was a putative prolyl 4-hydroxylase domain detected in this organism's proteome? TRUE = yes; FALSE = no. NA for non-leaf nodes.
'has_cthr': If this node is a leaf node, was a collagen domain detected in this organism's proteome? TRUE = yes; FALSE = no. NA for non-leaf nodes.
'max_cthr_length': If this node is a leaf node, the maximum collagen repeat length, in amino acids, of any protein detected in this organism's proteome. NA for non-leaf nodes. 0 for leaf nodes for which 'has_cthr' is FALSE.
'max_pavg': If this node is a leaf node, the maximum proline content of any protein detected in this organism's proteome. NA for non-leaf nodes. 0 for leaf nodes for which 'has_cthr' is FALSE.
'multiple_proteomes': If this node is a metazoan leaf node, are there multiple proteomes on UniProt for this node's Organism ID? TRUE = yes; FALSE = no. NA for non-metazoan nodes or non-leaf nodes.
'proteome_id': If this node is a metazoan leaf node, the UniProt Proteome ID for this node's Organism ID. If multiple_proteomes = TRUE for this node, then the proteome with the highest protein count is represented (note that this may not be the proteome in which the organism's collagen domain or prolyl 4-hydroxylase domain was found, if applicable). NA for non-metazoan nodes or non-leaf nodes.
'protein_count': If this node is a metazoan leaf node, the protein count corresponding to this row's Proteome ID. NA for non-metazoan nodes or non-leaf nodes.
'busco_c': If this node is a metazoan leaf node, the BUSCO-C score corresponding to this row's Proteome ID. NA for non-metazoan nodes or non-leaf nodes.
'cpd': If this node is a metazoan leaf node, the CPD corresponding to this row's Proteome ID. NA for non-metazoan nodes or non-leaf nodes.
'genrep': If this node is a metazoan leaf node, the Genome_Representation_Refseq corresponding to this row's Proteome ID. NA for non-metazoan nodes or non-leaf nodes.

3. SupplementaryDataFile3.csv

Description: Data for phylogenetic trees represented in Fig. 2C-F, Fig. S5A-B, Fig. S6. Each row represents a node of the phylogenetic tree.

'from': The parent taxon of the node that this row represents.
'to': The name of the node that this row represents. For leaf nodes, the name is exactly as found in the UniProt database.
'clade': The major eukaryotic group to which this node belongs (e.g., Archaeplastida, Discoba, SAR, etc.).
'family': The name of the family to which this node belongs. NA for nodes higher than family level.
'metazoan': Is this node within Metazoa? TRUE = yes; FALSE = no.
'isPhylum': If this node is within Metazoa, is it a phylum? TRUE = yes; FALSE = no. NA for non-metazoans.
'isFamily': Is this node a family? TRUE = yes; FALSE = no.
'isLeaf': Is this node a leaf node? TRUE = yes; FALSE = no.
'ox': The UniProt Organism ID corresponding to this node.
'has_ph': If this node is a leaf node, do any of the proteomes for this Organism ID encode a prolyl 4-hydroxylase domain? TRUE = yes; FALSE = no. NA for non-leaf nodes.
'has_cthr': If this node is a leaf node, do any of the proteomes for this Organism ID encode a collagen domain? TRUE = yes; FALSE = no. NA for non-leaf nodes.
'max_cthr_length': If this node is a leaf node, the maximum collagen repeat length, in amino acids, of any protein detected in any of the proteomes for this Organism ID. NA for non-leaf nodes. 0 for leaf nodes for which 'has_cthr' is FALSE.
'max_pavg': If this node is a leaf node, the maximum proline content of any collagen domain-containing protein detected in any of the proteomes for this node's Organism ID. NA for non-leaf nodes. 0 for leaf nodes for which 'has_cthr' is FALSE.
'multiple_proteomes': If this node is a leaf node, are there multiple proteomes on UniProt for this node's Organism ID? TRUE = yes; FALSE = no. NA for non-leaf nodes.
'proteome_id': If this node is a leaf node, the UniProt Proteome ID for this node's Organism ID. If multiple_proteomes = TRUE for this node, then the proteome with the highest protein count is represented. NA for non-leaf nodes.
'protein_count': If this node is a leaf node, the protein count corresponding to this row's Proteome ID. NA for non-leaf nodes.
'busco_c': If this node is a leaf node, the BUSCO-C score corresponding to this row's Proteome ID. NA for non-leaf nodes.
'cpd': If this node is a leaf node, the CPD corresponding to this row's Proteome ID. NA for non-leaf nodes.
'genrep': If this node is a leaf node, the Genome_Representation_Refseq corresponding to this row's Proteome ID. NA for non-leaf nodes.
'included': If this node is a leaf node, was this organism included in the collagen and prolyl 4-hydroxylase analyses? TRUE = yes; FALSE = no. NA for non-leaf nodes.
